# Neuroendocrine control of the proteostatic network by HPK-1 delays aging

**DOI:** 10.1101/2022.04.02.486836

**Authors:** Maria I. Lazaro-Pena, Carlos A. Diaz-Balzac, Ritika Das, Andrew V. Samuelson

## Abstract

The nervous system systemically coordinates proteostasis to delay organismal aging. However, the neuronal regulatory mechanisms that coordinate cellular anti-aging programs across tissue and cell-types are relatively unknown. In this work, we identify the <u>h</u>omeodomain-interacting <u>p</u>rotein <u>ki</u>nase (HPK-1), a transcriptional cofactor, as a novel neuronal component of the proteostatic network: its overexpression produces a paracrine signal to hyper-induce molecular chaperones and a neuroendocrine signal to induce autophagy in peripheral tissues. Neuronal HPK-1 signaling improves proteostasis in distal tissues through neurotransmitters. These pro-longevity modalities are independently regulated within serotonergic and GABAergic neurons, respectively, through distinct adaptive responses, either of which improve proteostasis in a cell non-autonomous manner. Serotonergic HPK-1 activity amplifies the heat shock response and protects the proteome from acute stress, without altering longevity. Conversely, increased GABAergic HPK-1 activity is sufficient to induce autophagy and extend longevity, without altering acute stress survival. Consistently, GABAergic neurons, but not serotonin, is essential for the cell non-autonomous induction of autophagy by neuronal HPK-1. These findings provide novel insight into how the nervous system partitions and coordinates unique adaptive response pathways to delay organismal aging, and reveals a key role for neuronal HPK-1 in regulating the proteostatic network throughout an intact metazoan animal.

**Significance Statement:** Aging and the age-associated decline of the proteome is determined in part through neuronal control of evolutionarily conserved transcriptional effectors, which safeguard homeostasis under fluctuating metabolic and stress conditions by regulating an expansive proteostatic network in peripheral tissues. How neuronal signaling mechanisms are primed, relayed through an organism, and specific responses are initiated in receiving cell types remain poorly understood. We have discovered that the *Caenorhabditis elegans* homeodomain-interacting protein kinase (HPK-1) is a novel transcriptional effector that functions within two distinct neuronal cell-types to non-autonomously regulate divergent components of the proteostatic network to enhance stress resistance, improve proteostasis and delay aging.

## Introduction

Cellular aging throughout an organism is characterized by the gradual decline of function within the proteome (proteostasis), which precipitates the onset and progression of a growing number of age-associated diseases both within and outside of the nervous system (1–9). A growing number of studies suggest that aging is not simply the result of the stochastic accumulation of damage, but is determined through genetics and coordinated mechanisms across tissues and cell types (10–12). This is not to say that aging or proteostatic decline occurs through a predetermined “clock”; rather genetic variation predisposes some individuals to age well and others to age poorly. Proteostatic decline corresponds with the age-associated breakdown of a large proteostasis network (PN), which coordinates stress-responsive control of protein folding, degradation, and translation in response to myriad cell intrinsic and non-autonomous signals. And yet, while major components and regulators of the proteostatic network have been identified, it is still poorly understood how multi-cellular organisms coordinate the maintenance of proteostasis across tissues to delay aging.

The relatively simple metazoan animal *Caenorhabditis elegans* (*C. elegans*) is a premiere model system for discovering how proteostasis is coordinated across cell- and tissue- types in response to myriad signals, and how these homeostatic processes breakdown during normal aging. The ability to examine complex cell non-autonomous regulatory interactions are derived from an invariant cell-lineage, a simplified nervous system, powerful tools to dissect cell- and tissue- type genetic interactions, a wealth of both genetic mutants and molecular/cellular reporters of the proteostatic network, and well characterized, quantifiable markers of age-associated decline in the proteome (12, 13). Discoveries of cell non-autonomous signaling first made in *C. elegans* are increasingly being demonstrated to have evolutionarily conserved components (14), which suggests overarching biological principles and mechanisms exist in how metazoans maintain proteostasis, and how they fail during normal aging.

An increasing number of studies have revealed the nervous system as a key cell non-autonomous regulator of organismal proteostasis and longevity in *C. elegans* (reviewed in (14)). For example, a pair of thermosensory neurons regulate the heat shock response (HSR) in distal tissue in a serotonin-dependent manner (15, 16). In contrast, a second regulatory component from the GABAergic and cholinergic system normally limits muscle cell proteostasis (17, 18). Interestingly, neuronal overexpression of heat shock transcription factor 1 (HSF-1) increases stress resistance and longevity through separable mechanisms (19). In accordance to these findings, another study found that integrin-linked kinase (ILK) inhibition activates HSF-1 cell non-autonomous effects on stress resistance and lifespan in a thermosensory-dependent manner (20). These studies highlight the key role of neuronal signaling in the regulation of proteostasis in distal tissues. Despite these advances, how neuronal cell non-autonomous signaling is activated under different adaptive cellular responses to metabolic and acute stress upon the proteome remains largely unknown.

In *C. elegans,* the transcriptional co-factor <u>h</u>omeodomain-interacting <u>p</u>rotein <u>ki</u>nase (HPK-1), preserves proteostasis and organismal longevity (21–23). We have previously shown that HPK-1 extends longevity and delays age-associated decline in proteostasis through distinct genetic pathways defined by the heat shock transcription factor (HSF-1), and the target of rapamycin complex 1 (TORC1) (23). Activation by either metabolic or genotoxic stressors has been observed from yeast to mammals, suggesting that this family of transcriptional cofactors arose early in evolution to couple metabolic and stress signaling (24–27).

The mammalian Hipk family consists of four conserved kinases (Hipk1-4), functioning as transcriptional co-factors that regulate cell growth, development, differentiation and apoptosis (28, 29). In general, HIPK family members regulate the activity of transcription factors, chromatin modifiers, signaling molecules and scaffolding proteins in response to cellular stress; including the DNA damage response, hypoxia response, reactive oxygen species (ROS), glucose availability, and viral infection (28, 30–32). For example, genotoxic damage induces mammalian *Hipk2,* and HIPK2 potentiates p53 pro-apoptotic activity through direct phosphorylation (24). Nutrient stress, such as glucose deprivation can also activate Hipk1, as well as Hipk2 (32–34). Conversely, hyperglycemia triggers HIPK2 degradation via the proteasome (35).

We sought to identify the tissue- and cell-types in which the HPK-1 transcriptional circuits acts, and how signals and regulatory components are coordinated and integrated across tissues to determine organismal longevity. We have discovered that HPK-1 functions as a key regulator of the proteostatic response, originating in the nervous system of *C. elegans*. HPK-1 acts in the nervous system to stimulate the release of both paracrine and endocrine signals that cell non-autonomously trigger protective peripheral responses. These responses can qualitatively differ depending on the neuronal type from which they arise. Serotonergic HPK-1 activity increases thermotolerance and protects the proteome from acute stress by hyper-induction of the heat shock response, without increasing lifespan or altering basal autophagy. In contrast, GABAergic HPK-1 activity fortifies the proteome by inducing autophagy to extend longevity, without altering thermotolerance. Yet, each of these distinct adaptive responses improves proteostasis in non-neuronal tissues. Our findings reveal HPK-1 as a novel neuronal regulator that coordinates adaptive metabolic and stress response pathways across tissues to delay aging throughout an animal.

## Results

### HPK-1 acts from neuronal tissue to promote longevity and health span

Organismal longevity is dependent upon the continued fidelity of a growing number of homeostatic mechanisms, which often have distinct spatial and temporal regulation. We sought to determine where HPK-1 exerts a regulatory control of longevity. We previously found that loss of *hpk-1* solely within either neuronal, intestinal, or hypodermal cells is sufficient to shorten lifespan (23). However, while *hpk-1* null-mutant animals are viable and fertile, HPK-1 and orthologs are known to have roles in development, differentiation and cell fate-specification (28, 29, 36–39). Thus, even though *hpk-1* null mutants display signs of premature aging and *hpk-1* overexpression throughout the soma extends longevity (23), it is possible that loss of *hpk-1* in some tissues may shorten adult lifespan by subtle developmental deficiencies independent of a role in aging.

To precisely determine where HPK-1 functions in the regulation of longevity, we overexpressed *hpk-1* in different tissues to identify where activity was sufficient to increase lifespan. We expressed *hpk-1* cDNA in either neuronal, muscle, intestine or hypodermis using well characterized tissue specific promoters (*rab-3p::hpk-1, myo-3p::hpk-1, ges-1p::hpk-1* and *dpy-7p::hpk-1*, respectively). Transgenic animals overexpressing *hpk-1* in either muscular, intestinal, or hypodermal cells had a normal lifespan (Fig 1B-D). In contrast, neuronal overexpression of *hpk-1* was sufficient to significantly increase lifespan by 17%, a modest amount comparable to constitutive overexpression of *hpk-1* throughout the soma (Fig. 1A) (23).

**Figure 1.**
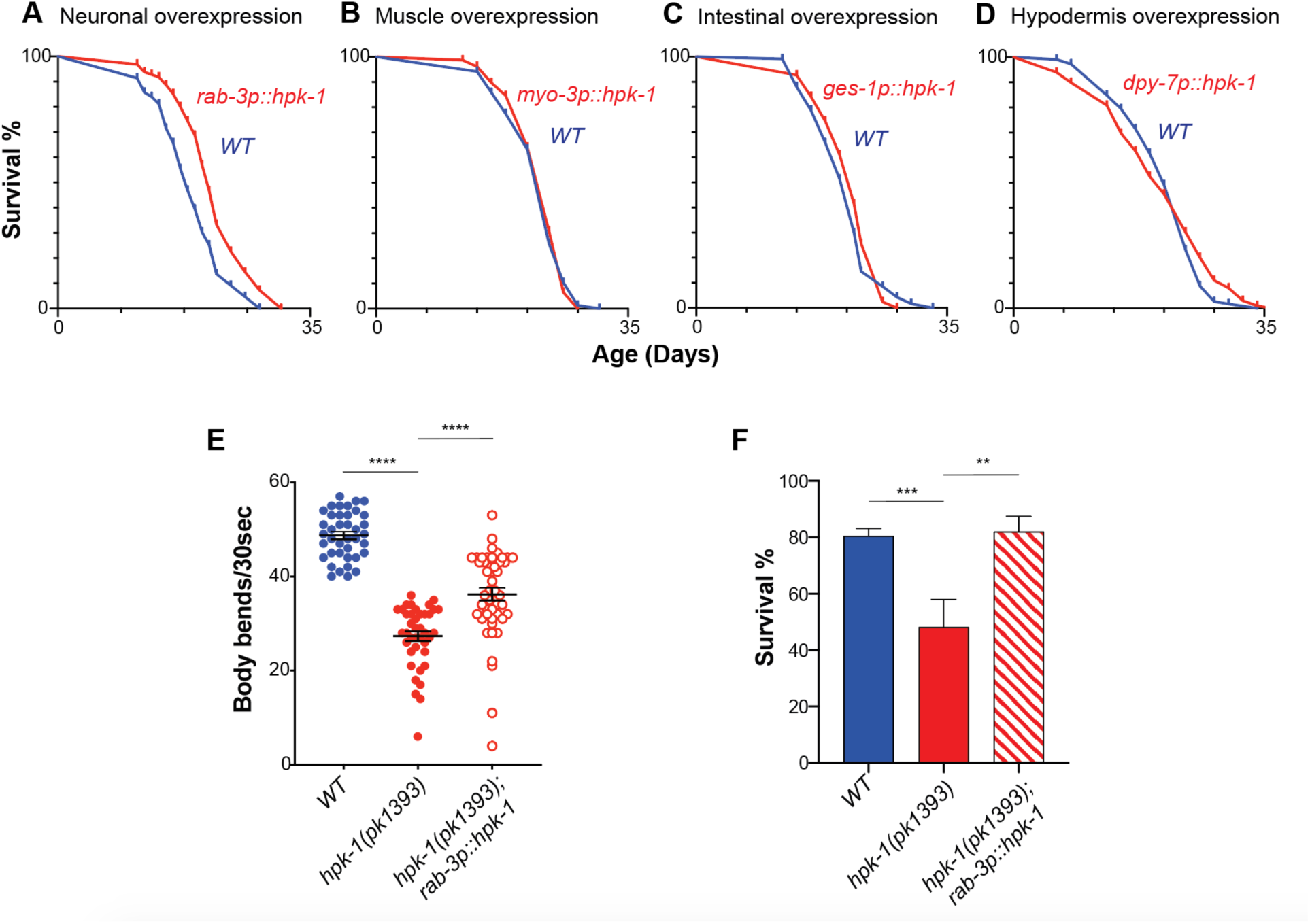
Neuronal HPK-1 extends longevity and promotes health span. Survival curves of animals overexpressing *hpk-1* (red line) in the nervous system **(A)**, muscle **(B)**, intestine **(C)** and hypodermis **(D)** compared with control non-transgenic siblings (blue line). (n≥175, *P* < 0.0001 for A, n≥78 for B, n≥82 for C and n≥105 for D). Lifespan graphs are representative of two biological replicates. **(E)** Frequency of body bends of *wild type*, *rab-3p::hpk-1* and *hpk-1(pk1393); rab-3p::hpk-1* day 2 adult animals maintained (n≥48). **(F)** Survival of day 1 adult animals subjected to heat shock treatment. Graph represents one of the two individual trials (n≥66). See Supplemental Tables 2, 4 and 6 for details and additional trials.

Because *hpk-1* null mutant animals are short-lived, and aging correlates with decreased movement (21-23, 40, 41), we performed a locomotion assay. *hpk-1(pk1393)* null mutants displayed a reduced number of body bends at day 2 of adulthood (Fig. 1E). However, overexpression of *hpk-1* in neurons was sufficient to mitigate the locomotion defect of *hpk-1* null mutants. We and others previously found that in the absence of *hpk-1*, animals are vulnerable to acute thermal stress (22, 23). Restoring *hpk-1* expression solely within the nervous system was sufficient to fully rescue the reduced thermotolerance of *hpk-1* null mutant day 1 adult animals (Fig. 1F). We conclude that *hpk-1* primarily acts within the nervous system to promote longevity, health span, and thermal stress resistance.

### Neuronal HPK-1 regulates muscle proteostasis via neurotransmission

The observed beneficial role of neuronal HPK-1 to extend longevity led us to postulate whether neuronal HPK-1 activates a cell extrinsic signal to regulate proteostasis in distal tissues. To investigate whether HPK-1 activity in neurons regulates distal muscle proteostasis, we used a proteostasis reporter expressed exclusively in muscle tissue (*unc-54p::Q35::YFP*) (42). Animals with increased neuronal *hpk-1* expression had significantly decreased fluorescent foci in muscle tissue (Fig. 2A*i-ii*). For instance, at day 3 of adulthood, animals overexpressing neuronal HPK-1 had an average of 21 ± 0.9 polyQ aggregates, while control animals had an average of 26 ± 1.1 polyQ aggregates (Fig. 2B). In *C. elegans*, the age-related progressive proteotoxic stress of muscle polyQ expression is pathological and results in paralysis. Of note, *hpk-1* overexpression in neurons was sufficient to reduce paralysis; at day 13 of adulthood only 61% ± 0.10 of neuronal *hpk-1* overexpressing animals were paralyzed compared to 81% ± 0.08 of paralyzed control animals (Fig. 2C).

**Figure 2.**
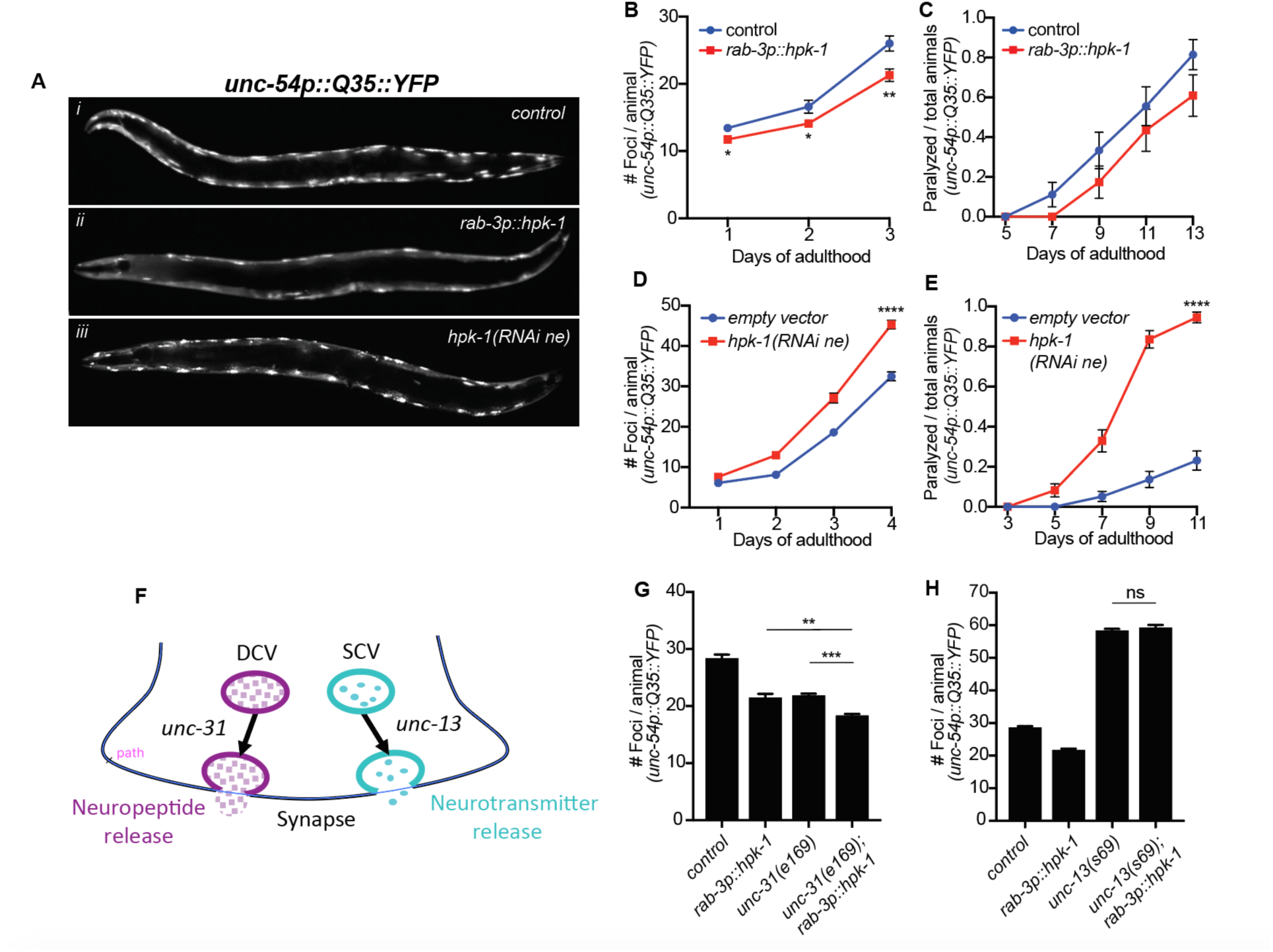
Neuronal HPK-1 prevents the decline in proteostasis through neurotransmitter release. **(A)** Fluorescent micrographs of animals expressing the polyglutamine fluorescent reporter in muscle. (**B** and **C**) Quantification of foci in muscle (B) and paralysis rate (C). Graphs are representative of 5 individual transgenic lines (n≥18 for B and n≥23 for C). (**D** and **E**) Quantification of foci in muscle (D) and paralysis rate (E) in neuronal enhanced RNAi animals (n≥22 for D and n≥73 for E). **(F)** Neuropeptides and neurotransmitters are essential for neuronal signaling; vesicle release depends on *unc-31* and *unc-13*, respectively. (**G** and **H**) Quantification of foci in muscle (G) (n≥21), (H) (n≥21). *T* test analysis with \*\**P* < 0.01 and \*\*\**P* < 0.001. See Supplemental Tables 3 and 5 for details and additional trials.

We next tested whether neuronal HPK-1 is required for maintaining muscle proteostasis. The nervous system of wild-type *C. elegans* is largely refractory to RNAi, as these cells do not express *sid-1*, which is necessary for dsRNA transport into neurons (43–45). Enhanced neuronal RNAi can be obtained by ectopic neuronal expression of *sid-1* (46). Neuronal inactivation of *hpk-1* (*sid-1(pk3321); unc-119p::sid-1; unc-54p::Q35::YFP; hpk-1(RNAi)*) was sufficient to hasten the collapse of proteostasis within muscle cells, significantly increasing the number of polyQ aggregates (45 ± 1 vs 32 ± 1.1 at day 4) and paralysis rate (84% ± 0.04 vs 14% ± 0.04 at day 9) (Fig. 2A*iii*,D,E). Thus, HPK-1 functions within the nervous system regulates proteostasis in distal muscle tissue, which implies that HPK-1 coordinates organismal health through cell non-autonomous mechanisms.

To identify the signaling mechanism employed by HPK-1 to regulate distal proteostasis, we tested whether mutant animals impaired for neuropeptide and neurotransmitter transmission are essential for cell non-autonomous regulation of proteostasis (*unc-31* and *unc-13* mutants, respectively Fig. 2F). Neuronal expression of *hpk-1* in *unc-31* mutants improved proteostasis in the absence of neuropeptide transmission (Fig. 2G). In contrast, in the absence of neurotransmitter function, neuronal expression of *hpk-1* in *unc-13* mutants failed to improve proteostasis in muscle tissue (Fig. 2H). Thus, HPK-1 neuronal cell non-autonomous signaling occurs through neurotransmitter transmission.

### Neuronal HPK-1 induces molecular chaperones and autophagy in distal tissues

We sought to identify the components of the PN that are cell non-autonomously regulated via HPK-1 activity in the nervous system. We first tested whether neuronal HPK-1 overexpression alters the expression of molecular chaperones (*hsp-16.2p::GFP*) and found basal expression of molecular chaperones was unchanged (Supplementary Fig. 1A). In contrast, after heat shock, neuronal *hpk-1* overexpression increased *hsp-16.2p::GFP* expression, particularly in the pharynx and head muscles, but surprisingly not in intestinal cells (Fig. 3A,B, Supplementary Fig. 1B,C). We conclude that while *hpk-1* is required for the broad induction of molecular chaperones, neuronal HPK-1 primes inducibility of the heat shock response (HSR) within specific tissues, rather than in a systemic manner throughout the organism.

**Figure 3.**
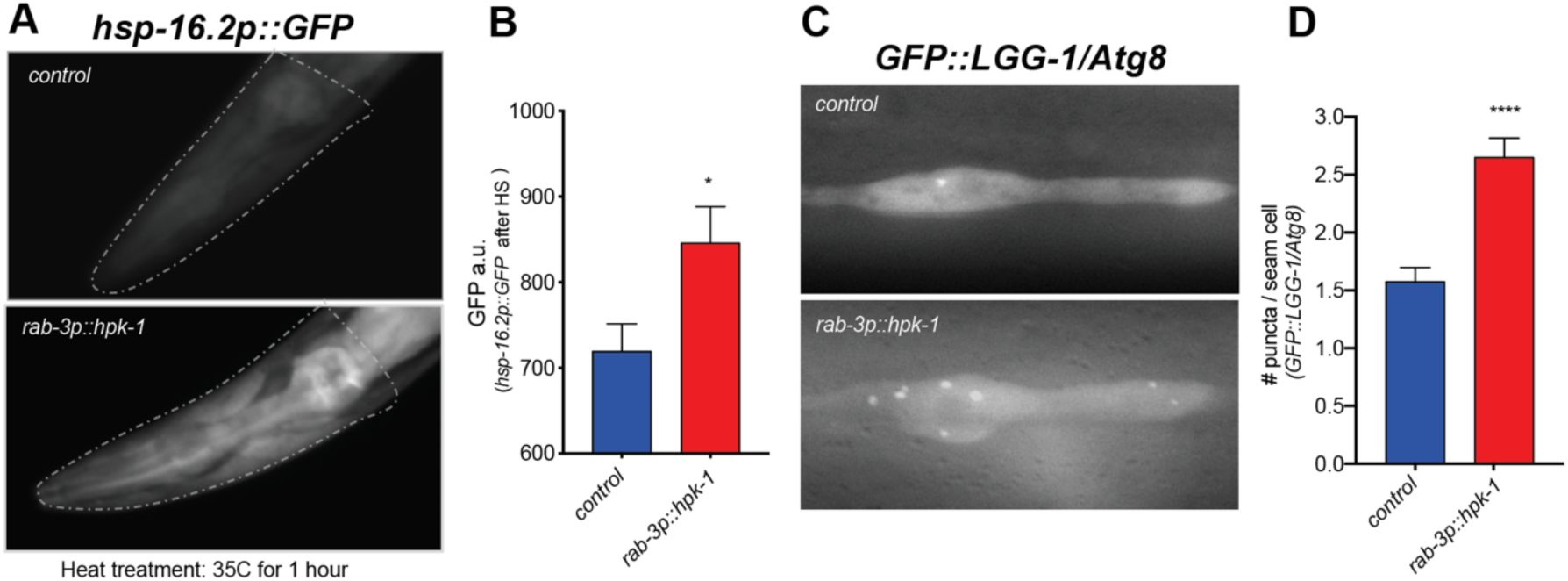
Neuronal HPK-1 induces the proteostatic network in distal tissues. **(A** and **B)** Fluorescent micrographs (A) and fluorescent density quantification (B) of control and *rab-3p::hpk-1* of day 1 adult animals expressing *hsp-16.2::GFP* molecular chaperone reporter that were exposed to heat shock for 1 hour at 35°C (n≥15). **(C** and **D)** Fluorescent micrographs (C) and quantification (D) of puncta in seam cells of control and *rab-3p::hpk-1* of L4 stage animals expressing the *lgg-1p::GFP::LGG-1/Atg8* autophagosome reporter (n≥43). *T* test analysis with \**P* < 0.05, \*\**P* < 0.01, \*\*\**P* < 0.001 and *\*\*\*\*P* < 0.0001. See Supplemental Tables 7 and 8 for details and additional trials.

We next tested whether increased neuronal HPK-1 activity is sufficient to induce autophagy. Using an LGG-1/Atg8 fluorescent reporter strain to visualize autophagosome levels in hypodermal seam cells (*lgg-1p::LGG-1::GFP)* (47), we found that neuronal *hpk-1* overexpression significantly induces autophagy (Fig. 3C,D), but not in the intestine (Supplementary Fig. 1D,E). Thus, increased HPK-1 activity within the nervous system acts to induce autophagy cell non-autonomously within a subset of peripheral tissues.

### Serotonergic and GABAergic HPK-1 differentially regulates heat stress response and autophagy

To begin to identify the specific neuronal cell-types from which neuronal HPK-1 initiates distinct cell non-autonomous signals, we overexpressed *hpk-1* in different subtypes of neurons and assessed changes in muscle proteostasis (*unc-54p::Q35::YFP*)*. hpk-1* overexpression in glutamatergic and dopaminergic neurons was not sufficient to decrease the formation of fluorescent foci or reduce paralysis (Supplementary Fig. 2). In contrast, overexpression of *hpk-1* in either serotonergic (*tph-1p::hpk-1*) or GABAergic (*unc-47p::hpk-1*) neurons was sufficient to improve proteostasis in muscle tissue and reduce paralysis (Fig. 4A-D). Accordingly, we found that *hpk-1* is expressed in serotonergic and GABAergic neurons, based on colocalization between *hpk-1* and neuronal cell-type specific reporters (Fig. 4E,F, Supplementary Fig. 3), which suggests that HPK-1 initiates pro-longevity signals from these neurons to preserve muscle proteostasis.

**Figure 4.**
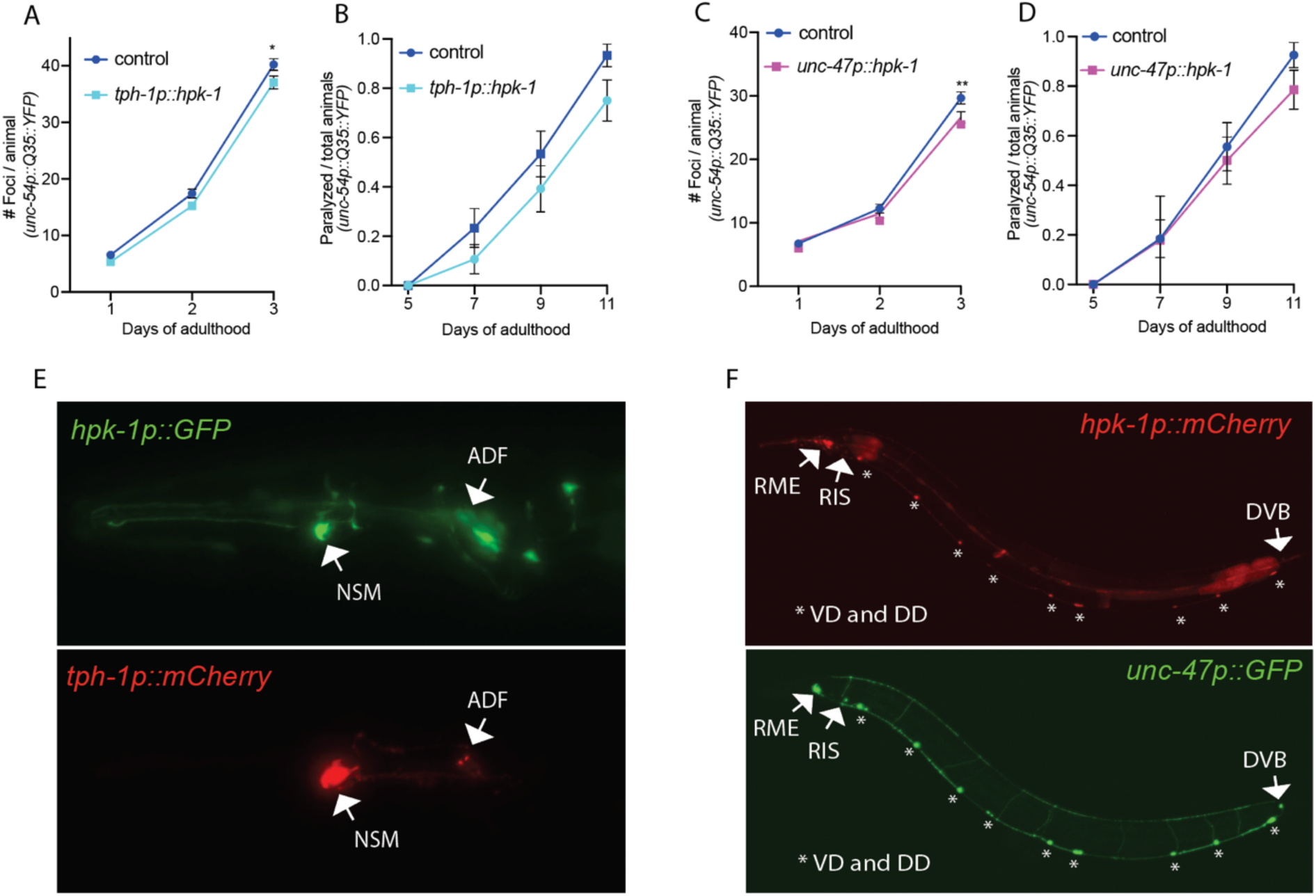
Expression of HPK-1 in serotonergic and GABAergic prevents the decline in proteostasis in muscle tissue. **(A** and **B)** Quantification of foci (A) and paralysis rate (B) of animals expressing *hpk-1* in serotonergic neurons (n≥17 for A and n≥28 for B). **(C** and **D)** Quantification of foci (C) and paralysis rate (D) of animals expressing *hpk-1* in GABAergic neurons (n≥19 for C and n≥27 for D). **(E)** Fluorescent micrographs of endogenous *hpk-1* expression in serotonergic neurons. **(F)** Fluorescent micrographs of endogenous *hpk-1* expression in GABAergic neurons. See Supplemental Tables: 3 and 5 for details and additional trials.

We sought to determine whether signals initiated from serotonergic and GABAergic neurons induced similar or distinct components of the PN in distal tissues, and assess the overall consequence on organismal health. Since pan-neuronal expression of HPK-1 was sufficient to induce autophagy, improve thermotolerance, and increase lifespan, we chose to focus on these three phenotypes. Increased HPK-1 serotonergic signaling was sufficient to improve thermotolerance, without altering autophagy or lifespan (Fig. 5A-C, Supplementary Fig. 4A,B). In contrast, increased HPK-1 expression in GABAergic neurons did not alter thermotolerance, but induced autophagy and slightly but significantly increased lifespan (Fig. D-F, Supplementary Fig. 4C,D).

**Figure 5.**
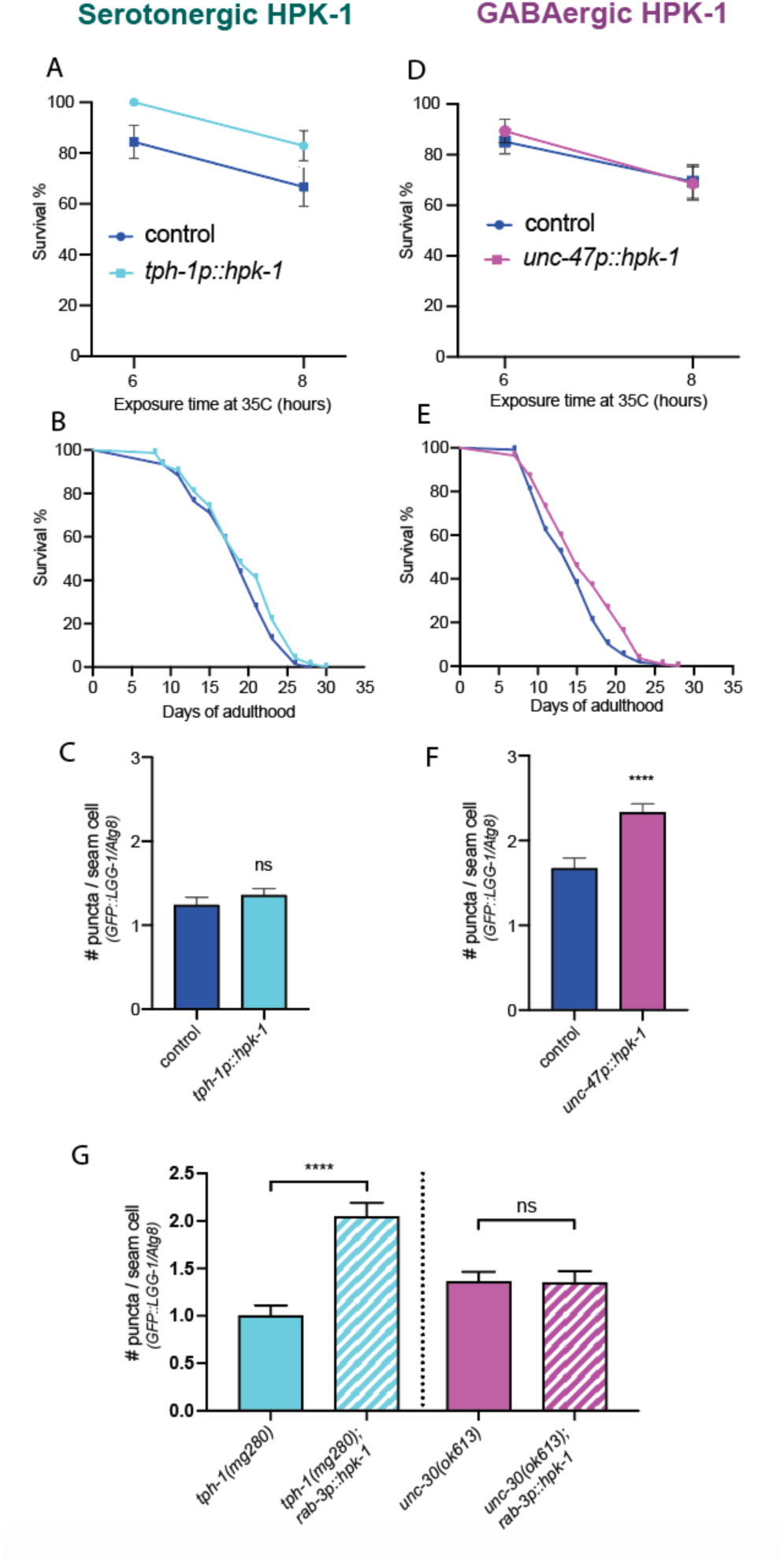
Serotonergic and GABAergic HPK-1 signaling activates distinct adaptive responses to regulate stress resistance and longevity. **(A)** Survival of serotonergic HPK-1 day 1 adult animals after heat shock (n≥32). **(B)** Lifespan of serotonergic HPK-1 (n≥76). **(C)** Quantification of autophagosomes in hypodermal cells of serotonergic HPK-1 day 1 adult animals (n≥80). **(D)** Survival of GABAergic HPK-1 day 1 adult animals after heat shock (n≥47). **(E)** Lifespan of GABAergic HPK-1 animals (n≥119). **(F)** Quantification of autophagosomes in hypodermal cells of GABAergic HPK-1 day 1 adult animals (n≥64). (**G**) Quantification of autophagosomes in hypodermal cells of mutant animals of serotonin and GABAergic signaling (*tph-1* and *unc-30*, respectively), expressing pan-neuronal HPK-1 (day 1 adults, n≥ 67). *T* test analysis with \**P* < 0.05, \*\**P* < 0.01 and *\*\*\*\*P* < 0.0001. See Supplemental Tables: 2, 6 and 8 for details and additional trials.

We sought to identify the mechanism through which increased neuronal HPK-1 activity induces autophagy in distal tissues. As expected loss of *tph-1,* which encodes the ortholog of human tryptophan hydroxylase 1 and is essential for the biosynthesis of serotonin (48), failed to block the induction of autophagy by increased neuronal *hpk-1* activity (Fig. 5G, *tph-1(mg280);rab-3p::hpk-1*)*. unc-30* encodes the ortholog of human paired like homeodomain 2 (PITX2) and is essential for the proper differentiation of type-D inhibitory GABAergic motor neurons (49, 50). Loss of *unc-30* was sufficient to completely abrogate the induction of autophagy in seam cells after pan-neuronal overexpression of HPK-1 (Fig. 5G, *unc-30(ok613);rab-3p::hpk-1*). Collectively, we conclude that HPK-1 initiates distinct signals from serotonergic and GABAergic neurons to improve organismal health. Specifically, HPK-1 initiates serotonergic signals to improve survival to acute thermal stress, likely as a part of the HSF-1 thermosensory circuit (discussed below), and regulates GABAergic signals to increase lifespan through autophagy in the absence of stress (Fig. 6).

**Figure 6.**
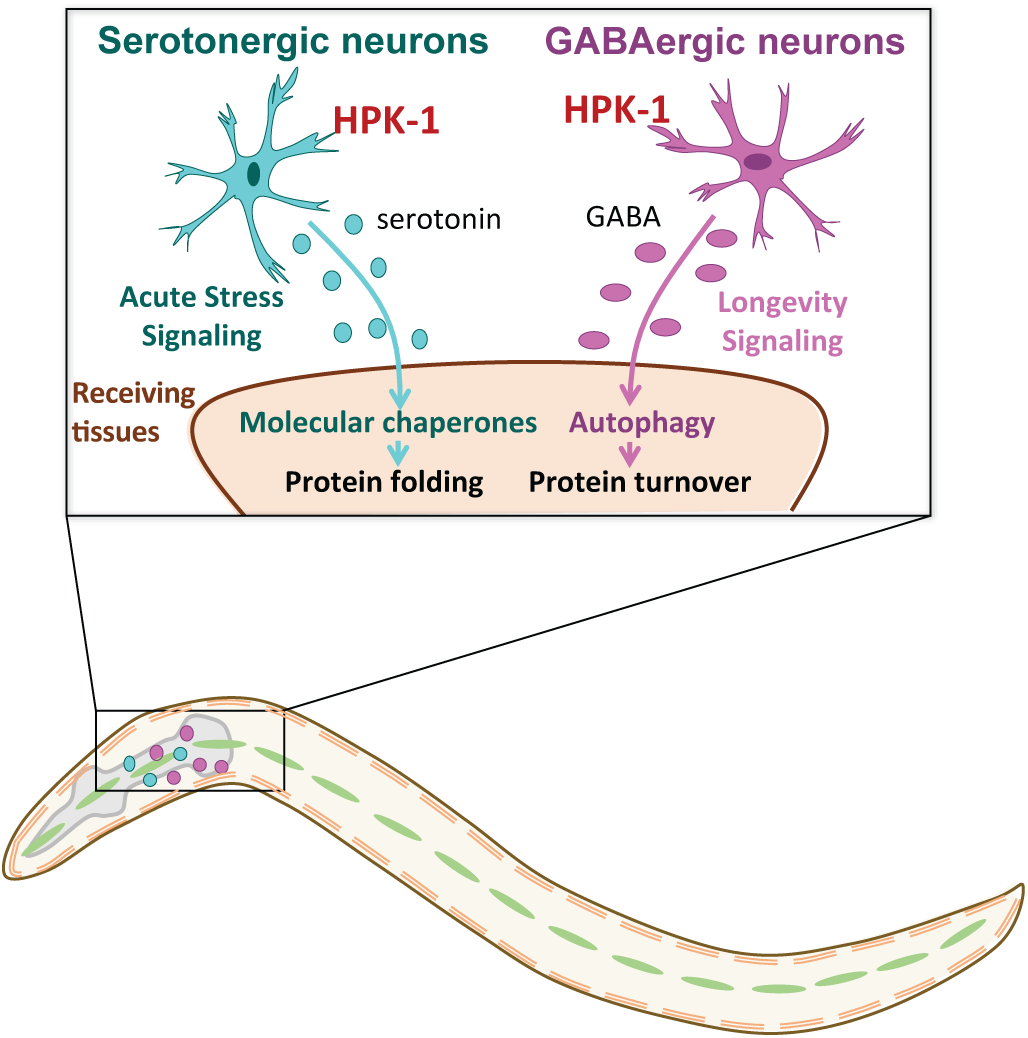
Model diagram of serotonergic and GABAergic HPK-1 regulatory mechanism to mediate proteostasis and longevity. HPK-1 activity in serotonergic and GABAergic neurons initiates distinct adaptive responses, either of which improve proteostasis in a cell non-autonomous manner. Serotonergic HPK-1 protects the proteome from acute stress, while GABAergic HPK-1 fortifies the proteome by regulating autophagy activity in response to other types of stress, such as metabolic stress.

## Discussion

In this study we show that HPK-1 expression in serotonergic neurons activates a cell non-autonomous signal to induce heat stress response and protein folding in muscles, whereas HPK-1 expression in GABAergic neurons signals the induction of autophagy activity in hypodermal tissue. A growing number of studies have demonstrated the nervous system plays a pivotable role in the regulation of overall organismal health and longevity. In response to metabolic, environmental, and intrinsic cues, the *C. elegans* nervous system can initiate a number of distinct stress and longevity signals cell non-autonomously, which activates comparable changes in gene expression peripheral tissues (15, 16, 18, 19, 51–55). We demonstrate a neuronal cell-type specific role for serotoninergic and GABAergic cell non-autonomous regulation of the PN by HPK-1. Loss of serotonin production in *tph-1* mutant animals has a minimal impact on lifespan at basal temperature (56). However, serotonin signaling is required for several conditions that extend longevity. For instance, animals treated with an atypical antidepressant serotonin antagonist (mianserin) (57), changes in food perception (58), and neuronal cell non-autonomous induction of the mitochondrial unfolded protein response (59, 60), all require *tph-1* and serotonin signaling. Furthermore, loss of the serotonin receptor *ser-1* is sufficient to increase lifespan (56). This is consistent with the notion that serotonergic neurons integrate diverse signals from sensory neurons to regulate gene expression in peripheral tissues that influence organismal longevity.

Surprisingly, we find that increased HPK-1 activity in serotonergic neurons does not increase lifespan, but is sufficient to improve muscle proteostasis and amplify the expression of molecular chaperones that maintain proteostasis within the cytosol and nucleus. Previous findings have discovered that ADF serotoninergic neurons receive upstream stress signals from the thermosensory circuit, consisting of the amphid sensory neurons (AFD) and the AIY interneurons, which regulate HSF-1 activity in the germline in a serotonin-dependent manner (16). Serotonin signaling and the serotonin receptor *ser-1* are essential for the induction of the HSR (15, 16). Interestingly, our results are analogous with a previous finding that demonstrated HSF-1 regulation of the HSR through thermosensory circuits is separable from lifespan (19). HPK-1 co-localizes with HSF-1 in neurons and is also essential for induction of the HSR (19, 23). Collectively, our results suggest that HPK-1 is an integral part of the thermosensory-serotonin circuit to maintain homeostasis in response to acute thermal stress.

Increased HPK-1 activity in GABAergic neurons induces autophagy in peripheral cells, improves proteostasis and longevity. Whether HPK-1 functions in GABAergic neurons through alteration of GABA or another neurotransmitter is unknown, but proper differentiation of GABAergic neurons is essential for neuronal HPK-1 activation of autophagy in distal tissue. We posit HPK-1 cell non-autonomous signaling occurs via antagonism of GABA signaling. Loss of GABA increases *C. elegans* lifespan (61, 62). Interestingly, in mice GABA signaling inhibits autophagy and causes oxidative stress, which is mitigated via Tor1 inhibition (63). In fact, neuronal expression of HPK-1 is inhibited by TORC1 and *hpk-1* is essential for TORC1 inactivation to extend longevity and induce autophagy gene expression (23). Of note, TORC1 acts within neurons to regulate aging (64). Collectively, our results position HPK-1 activity in GABAergic neurons as the likely integration point for cell non-autonomous regulation of autophagy via TORC1.

This work reveals a new level of specificity in the capacity of the nervous system to initiate adaptive responses by triggering distinct components of the proteostatic network across cell types. Elucidating how neuronal cell-types generate heterotypic signals to coordinate the maintenance of homeostasis are just beginning to emerge. For example, a previous study discovered that activating non-autonomous signaling of the endoplasmic reticulum unfolded protein response (ER-UPR) via the XBP-1 transcription factor can be initiated from serotonergic neurons and dopaminergic neurons; these signals are unique and converge to activate distinct branches of the ER-UPR within intestinal cells (65). In contrast, our work demonstrates differential, yet specific, activation of cytosolic components of the PN in separate peripheral tissues. Our work suggests an emerging biological principle for the cell non-autonomous regulation of longevity: the nervous system partitions neuronal cell-types to coordinate the activation of complementary proteostatic mechanisms throughout tissues, despite utilizing the same transcriptional regulator within each neuronal cell-type. This is consistent with recent findings: during development and differentiation, transcriptional rewiring of cells set chaperoning capacities and alter usage of mechanisms that maintain protein quality control (66, 67). Our work has revealed a novel insight into how an emerging transcriptional circuit within the nervous system of metazoans integrates diverse stimuli to coordinate specific adaptive responses to determine organismal aging.

## Materials and Methods

### *C. elegans* strains and details

All strains were maintained at 20°C on standard NGM plates with OP50. For all experiments, animals were grown in 20x concentrated HT115 bacteria seeded on 6cm RNAi plates. Details on the strains, mutant alleles, and transgenic animals used in this study are listed in Supplemental Table 1.

### Generation of transgenic strains

To assemble tissue-specific constructs for overexpression experiments, the *hpk-1* cDNA was cloned under control of the following promoters: hypodermal *dpy-7p* (*68*), body wall muscle *myo-3p* (*69*), pan-neuronal *rab-3p* (70), dopaminergic neurons *cat-2p* (*71*), glutamatergic neurons *eat-4p* (*72*), serotonergic neurons *tph-1p* (*48*), and g-aminobutyric acid (GABA)ergic neurons *unc-47p* (*73*). All plasmids contained the *unc-54* 3’-UTR. These constructs for were injected at 5 ng/ml together with *myo-2p::mCherry* construct at 5 ng/ml as injection marker.

### RNAi feeding

The *hpk-1* RNAi clone was originated from early copies of *E*. *coli* glycerol stocks of the comprehensive RNAi librariy generated in the Ahringer laboratory. The control empty vector (L4440) and *hpk-1* RNAi colonies were grown overnight in Luria broth with 50 μg ml–1 ampicillin and then seeded onto 6cm RNAi agar plates containing 5 mM isopropylthiogalactoside (IPTG) to induce dsRNA expression overnight at room temperature. RNAi clones used in this study were verified by DNA sequencing and subsequent BLAST analysis to confirm their identity.

The transmembrane protein *sid-1* is essential for dsRNA transport into the cell (43–45), however, *C. elegans* nervous system does not express *sid-1.* Enhanced neuronal RNAi can be obtained by ectopic neuronal expression of *sid-1* (46). Thus, we crossed the *rgef-1p::Q40::YFP* reporter with *sid-1(pk3321);unc-119p::sid-*1 for neuronal downregulation experiments.

### Lifespan analysis

Lifespan assays were performed essentially as described in: (23, 74). Briefly, animals were synchronized by egg prep bleaching followed by hatching in M9 solution at 20°C overnight. The L1 animals were seeded onto 6cm plates with HT115 bacteria and allowed to develop at 20°C. At L4 stage, 2’ fluoro– 5’deoxyuridine (FUDR) was added to a final concentration of 400 uM to prevent progeny production (defined as day 0 adulthood). Viability was scored every day or every other day as indicated in each figure. Prism 7 was used to for statistical analyses and the *P* values were calculated using log-rank (Mantel-Cox) method.

### Thermotolerance assay

Survival assays at high temperature were conducted as previously described in (75). In brief, synchronized L1 animals were allowed to develop at 20°C; FUDR was added at the L4 stage to prevent progeny production. At day 1 adulthood, animals were moved to 35°C for a period of 6 and 8 hours. Animals were allowed to recover for 2 hours at 20°C, and viability was scored. Statistical testing between pairs of conditions for differences in the number of foci was performed using student’s *t* test analysis.

### Induction of *hsp-16*.*2p*::*GFP* after heat shock

The *hsp-16*.*2p*::*GFP* animals were heat shocked on day 1 of adulthood for 1 hour at 35°C and imaged after 4 hours of recovery. Images were acquired using a Zeiss Axio Imager M2m microscope with AxioVision v4.8.2.0 software. The GFP a.u. values from the head and intestine of the animals were acquired using the area selection feature from AxioVision software. Two independent trials with a minimum of 20 animals per experiment were performed.

### Measurement of autophagosome formation using the *GFP::LGG-1/Atg8* reporter

*GFP::LGG-1/Atg8* foci formation was visualized as described in (47). Briefly, L4 stage and day 1 adult animals were raised on HT115 bacteria at 20°C and imaged using a Zeiss Axio Imager M2m microscope with AxioVision v4.8.2.0 software at 63X magnification. More than 40 seam cells from 15-20 different animals were scored for *GFP::LGG-1* puncta accumulation. Two independent trials with a minimum of 20 animals per experiment were performed.

### Polyglutamine aggregation in muscle and paralysis analyses

The visualization and quantification of the progressive decline in proteostasis in muscle tissue was performed as described in (42, 76). Briefly, synchronized *unc-54p*::*polyQ35*::*YFP* L4 animals were treated with FUDR to prevent progeny production. Then, fluorescent foci from 20 animals per technical replicate were scored blind daily from days one to three of adulthood. To assess paralysis, at days 5,7,9 and 11 of adulthood, prodded animals that responded with head movement (and were therefore still alive) but no backward or repulsive motion were scored as paralyzed (as described in (76)). Statistical testing between pairs of conditions for differences in the number of foci was performed using student’s *t* test analysis. Two to five independent transgenics lines were tested.

### Statistical analyses

Graphpad PRISM7 software was used for all statistical analyses. Data were considered statistically significant when *P* value was lower than 0.05. In figures, asterisks denote statistical significance as (*, *P* < 0.05, **, *P* < 0.001, ****, *P* < 0.0001) as compared to appropriate controls. N number listed in figure legend indicates the number of one representative experiment among all biological trials.

## Data availability

All data supporting the findings of this study are available within the article, its Supplementary Information files, the peer-review file, or are available upon request.

## Acknowledgments

We would like to thank members of the Samuelson laboratory, past and present, for their thoughtful insight in many discussions related to this project. In this regard, we would particularly like to thank Adam Cornwell. We would like to thank members of the Western New York Worm Group for their input and discussions. We would like to thank Dr. Doug Portman (URMC) and his laboratory for strains and advice. Some strains were provided by the CGC, which is funded by NIH Office of Research Infrastructure Programs (P40 OD010440). We would like to personally thank Dr. Dirk Bohmann (URMC) for the critical reading of this manuscript. Research reported in this publication was supported by the National Institute On Aging of the National Institutes of Health under Award Number RF1AG062593. CADB was supported by F32HD105323. The content is solely the responsibility of the authors and does not necessarily represent the official views of the National Institutes of Health.

## Supplementary Information

### Supplementary figures and figure legends

**Figure S1.**
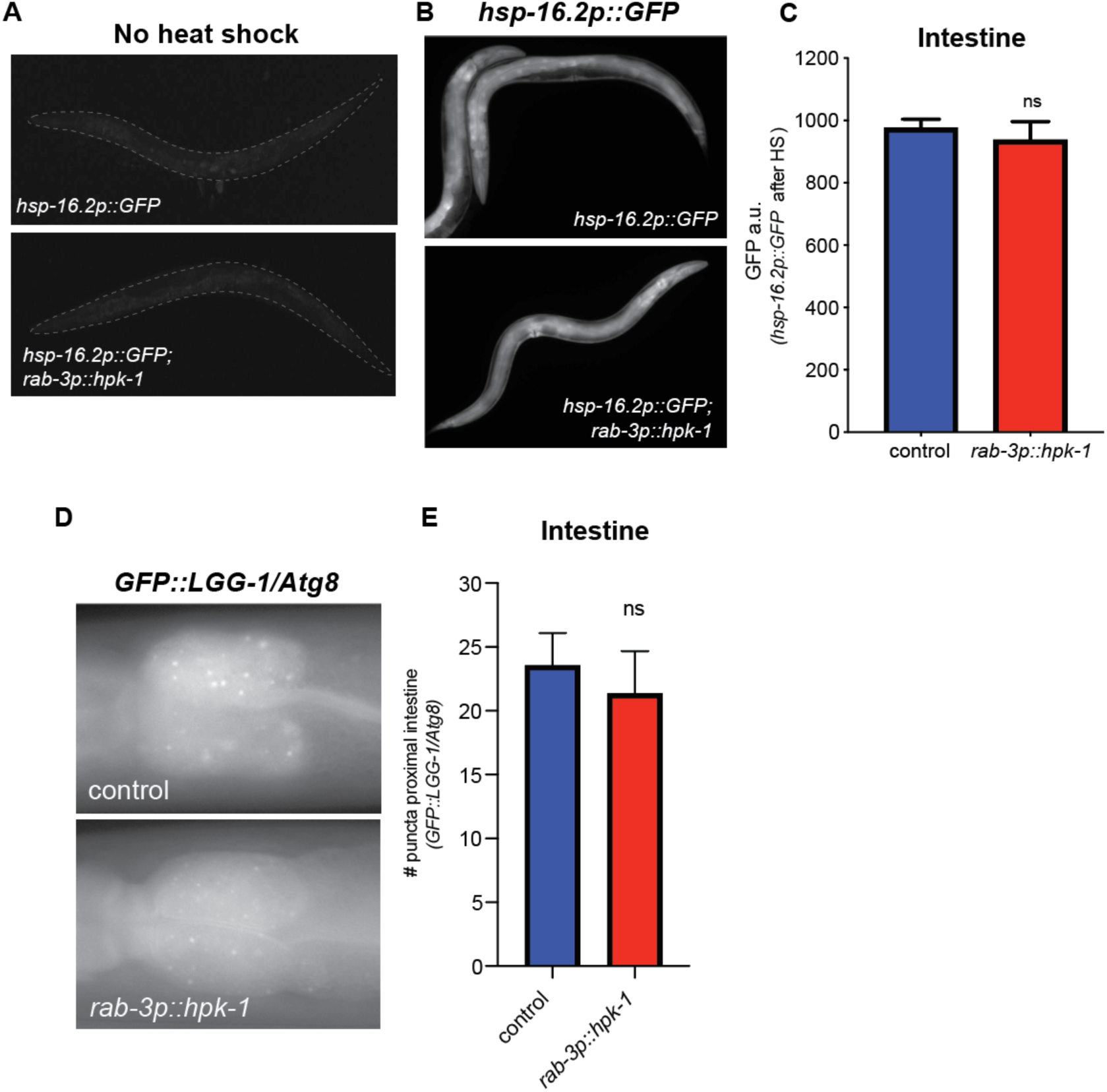
Neuronal HPK-1 does not hyper-induces molecular chaperones in the intestine. **(A**) Animals expressing *hsp-16.2p::GFP* in basal condition. **(B)** Animals expressing *hsp-16.2p::GFP* after heat shock treatment for 1hr at 35°C. **(C)** GFP densitometry quantification on the intestine of day 1 adults after heat treatment from *hsp-16.2p::GFP* expressing animals (n≥24). (D) Fluorescent micrographs and (E) quantification of puncta in proximal intestinal cells of control and *rab-3p::hpk-1* of L4 stage animals expressing the *lgg-1p::GFP::LGG-1/Atg8* autophagosome reporter (n≥17). See Supplemental Table 7 and 8 for details and additional trials.

**Figure S2.**
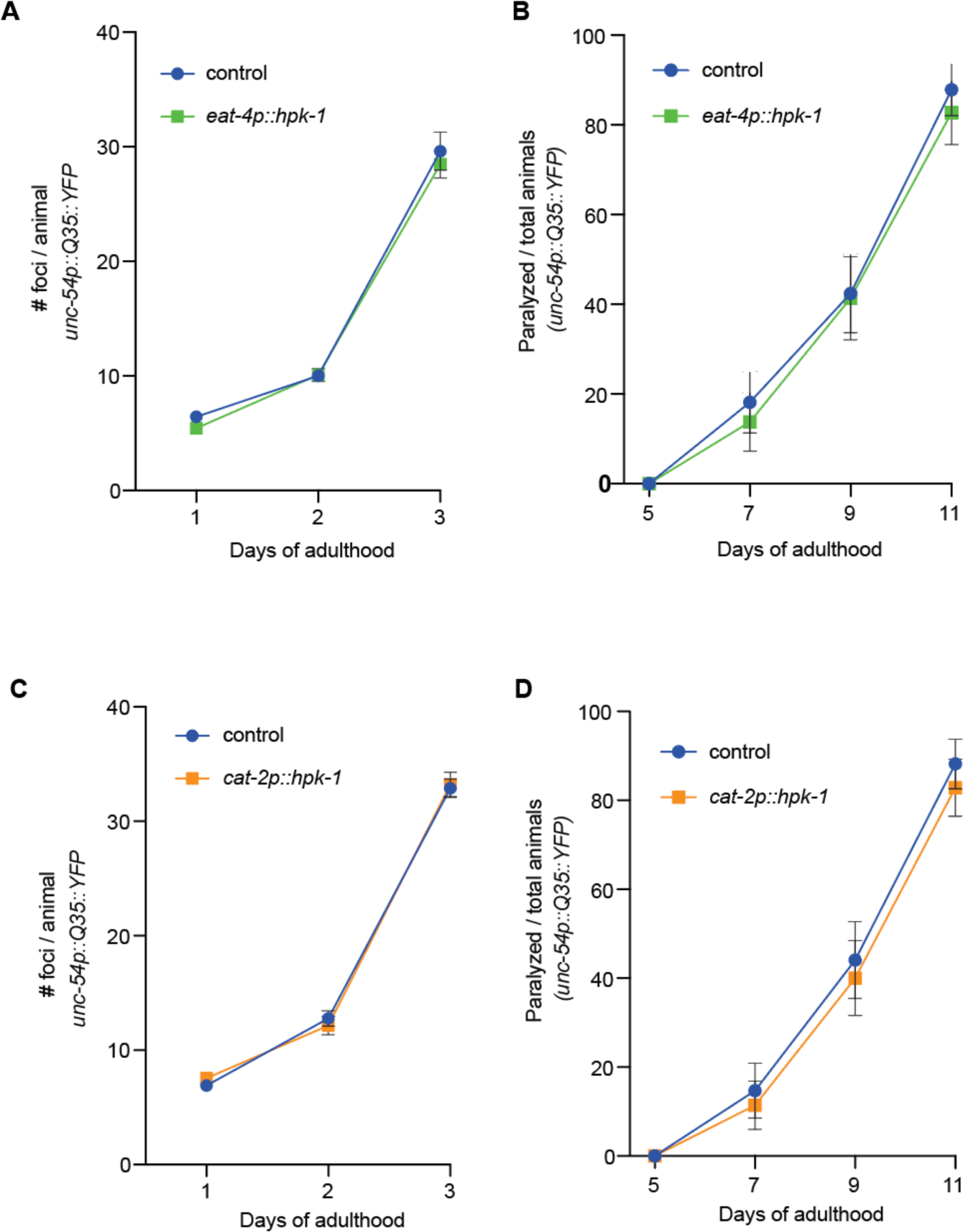
*hpk-1* expression in glutamatergic and dopaminergic neurons is not sufficient to prevent the decline in proteostasis. **(A** and **B)** Quantification of number of foci (A) and paralysis rate (B) of animals expressing *hpk-1* in the glutamatergic neurons (*eat-4p::hpk-1*; green line) and control non-transgenic siblings (blue line) (n≥18 for A and n≥29 for B). **(C** and **D)** Quantification of number of foci (C) and paralysis rate (D) of animals expressing *hpk-1* in the dopaminergic neurons (*cat-2p::hpk-1;* orange line) and control non-transgenic siblings (blue line) (n≥16 for C and n≥34 for D). See Supplemental Tables 3 and 5 for details and additional trials.

**Figure S3.**
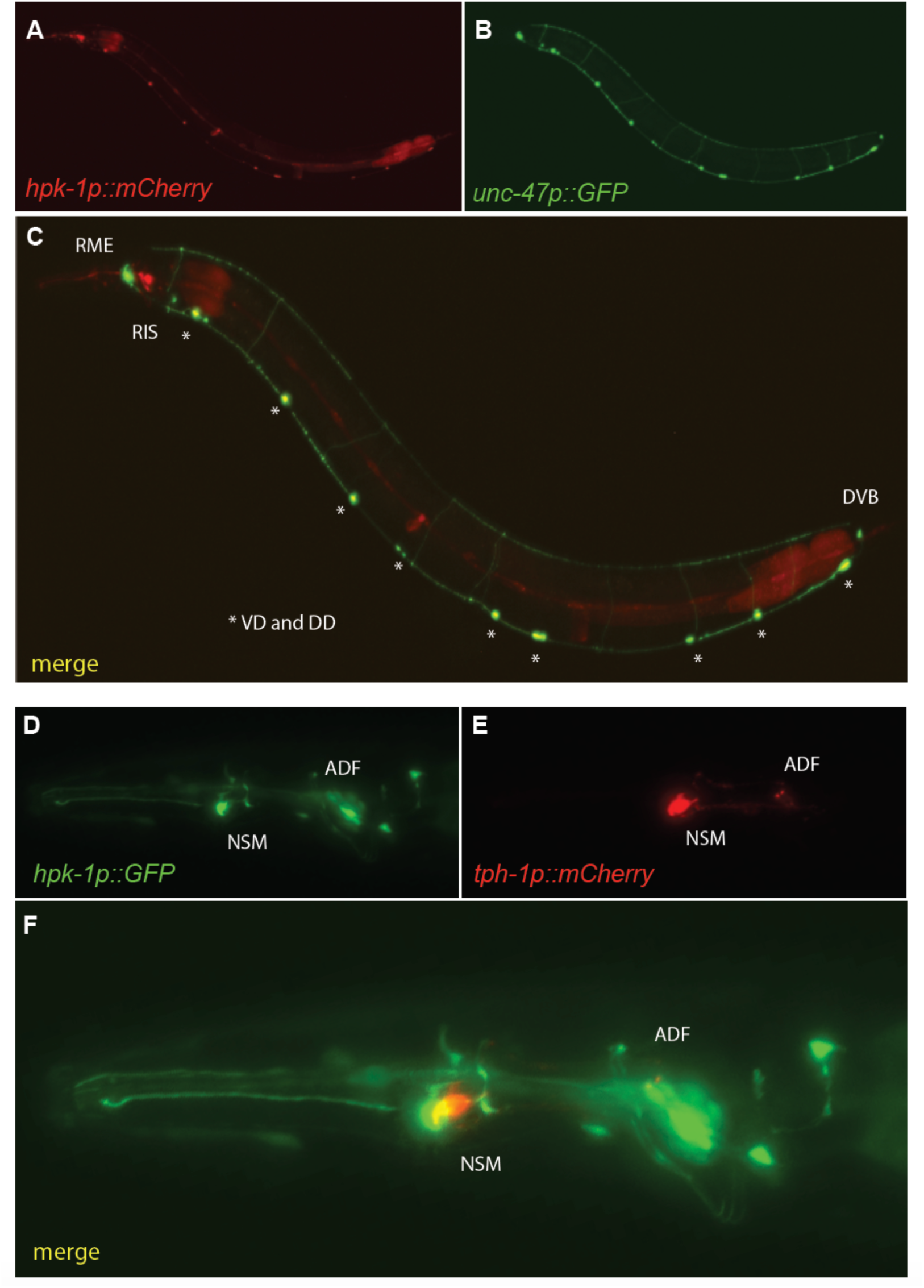
*hpk-1* is expressed in GABAergic and serotonergic neurons. Representative fluorescent micrographs of day 1 adult animals co-expressing *hpk-1p::mCherry* and *unc-47p::GFP* **(A-C)**; and *hpk-1p::GFP* and *tph-1p::mCherry* **(D-F)**.

**Figure S4.**
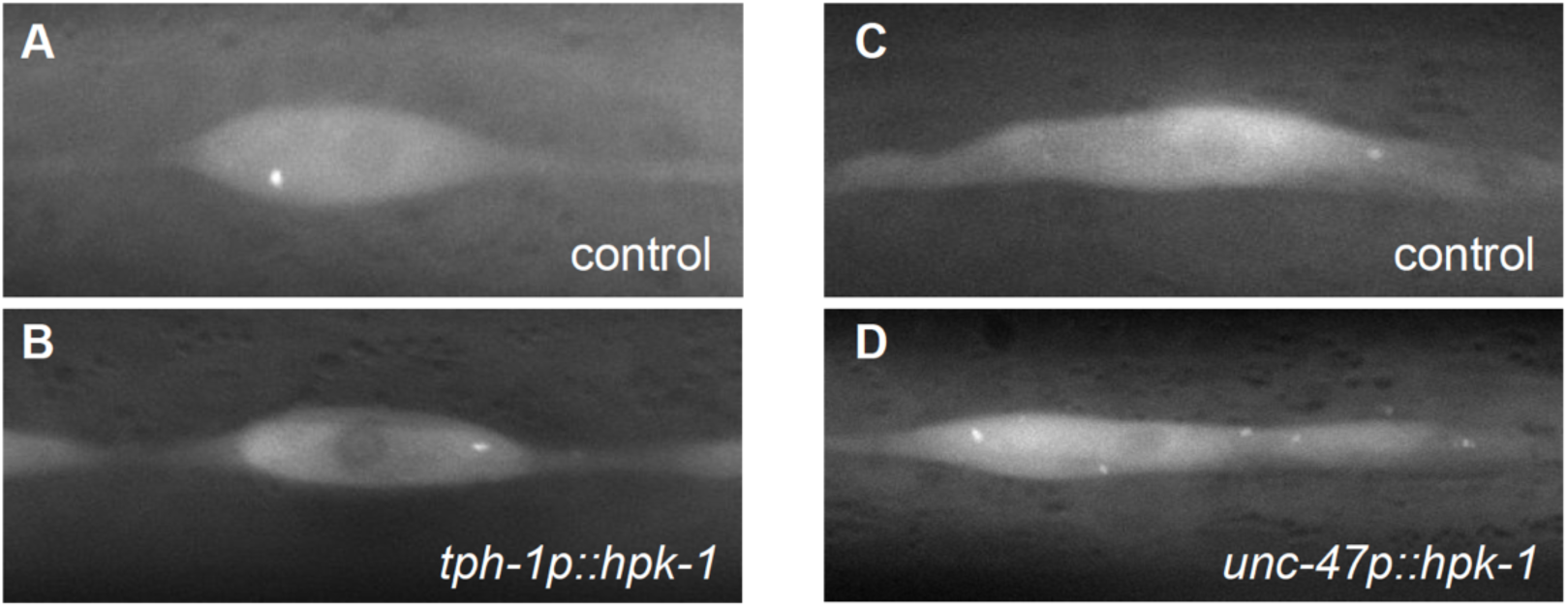
*hpk-1* expression in GABAergic neurons, but not in serotonergic neurons, induces autophagy activity in hypodermal seam cells. Representative fluorescent micrographs (from data in Figure 5I,K), showing autophagosome fluorescent puncta in seam cells of day 1 adult animals expressing *GFP::LGG-1/Atg8* autophagosome reporter with *hpk-1* expression in serotonergic (**A** and **B**) and GABAergic (**C** and **D**) neurons.

### Supplementary tables

**Table S1.** Strains, mutant alleles, and transgenic animals used in this study.

| Strain name | Genotype |
| --- | --- |
| N2 | *Wild type* |
| AVS392 | *hpk-1(pk1393) X* |
| AVS543 | *artEx41 [rab-3p::hpk-1::CFP + pCFJ90 (myo-2p::mCherry)]* |
| AVS609 | *artEx43 [myo-3p::hpk-1 + pCFJ90 (myo-2p::mCherry)] line 1* |
| AVS614 | *artEx48 [dpy-7p::hpk-1 + pCFJ90 (myo-2p::mCherry)] line 1* |
| AVS627 | *artEx52 [ges-1p::hpk-1 + pCFJ90 (myo-2p::mCherry)] line 1* |
| AVS420 | *hpk-1(pk1393) X; artEx41 [rab-3p::hpk-1::CFP + pCFJ90 (myo-2p::mCherry)]* |
| AVS832 | *rmls132 [unc-54p::Q35::YFP] I; artEx58 [rab-3p::hpk-1 + pCFJ90 (myo-2p::mCherry)] line 1* |
| AVS833 | *rmls132 [unc-54p::Q35::YFP] I; artEx59 [rab-3p::hpk-1 + pCFJ90 (myo-2p::mCherry)] line 2* |
| AVS834 | *rmls132 [unc-54p::Q35::YFP] I; artEx60 [rab-3p::hpk-1 + pCFJ90 (myo-2p::mCherry)] line 3* |
| AVS835 | *rmls132 [unc-54p::Q35::YFP] I; artEx61 [rab-3p::hpk-1 + pCFJ90 (myo-2p::mCherry)] line 4* |
| AVS562 | *rmls132 [unc-54p::Q35::YFP] I; artEx41 [rab-3p::hpk-1::CFP + pCFJ90 (myo-2p::mCherry)] line 5* |
| AVS214/HC196 | *sid-1(qt9) V* |
| AVS265/TU3401 | *sid-1(pk3321)V; uls69 [unc-119p::sid-1; myo-2p::mCherry]V* |
| AVS540 | *rmls132 [unc-54p::Q35::YFP] I; sid-1(pk3321)V; uls69[unc-119p::sid-1; myo-2p::mCherry]V* |
| AVS713 | *rmls132 [unc-54p::Q35::YFP] I; artEx41(rab-3p::hpk-1::CFP + pCFJ90 (myo-2p::mCherry)); unc-31(e169)V* |
| AVS810 | *rmls132 [unc-54p::Q35::YFP] I; artEx41 [rab-3p::hpk-1::CFP + pCFJ90 (myo-2p::mCherry)]; unc-13(e51)I* |
| AVS84/TJ375 | *gpls1 [hsp-16.2p::GFP] IV* |
| AVS397 | *gpls1 [hsp-16.2p::GFP]; artEx35 [sur-5p::hpk-1::CFP + pCFJ90 (myo-2p::mCherry)]* |
| AVS399 | *gpls1 [hsp-16.2p::GFP]; artEx33 [rab-3p::hpk-1::CFP + pCFJ90 (myo-2p::mCherry)]* |
| AVS716 | *artEx41 [rab-3p::hpk-1::CFP + pCFJ90 (myo-2p::mCherry)]; sqs13 [lgg-1p::GFP::lgg-1 + odr-1p::RFP]* |
| AVS682 | *artEx62 [tph-1p::hpk-1 + pCFJ90 (myo-2p::mCherry)] line 1; rmls132 [unc-54p::Q35::YFP] I* |
| AVS683 | *artEx63 [tph-1p::hpk-1 + pCFJ90 (myo-2p::mCherry)] line 2; rmls132 [unc-54p::Q35::YFP] I* |
| AVS685 | *artEx65 [unc-47p::hpk-1 + pCFJ90 (myo-2p::mCherry)] line 1; rmls132 [unc-54p::Q35::YFP] I* |
| AVS686 | *artEx66 [unc-47p::hpk-1 + pCFJ90 (myo-2p::mCherry)] line 2; rmls132 [unc-54p::Q35::YFP] I* |
| AVS691 | *artEx71 [eat-4p::hpk-1 + pCFJ90 (myo-2p::mCherry)] line 1; rmls132 [unc-54p::Q35::YFP] I* |
| AVS692 | *artEx72 [eat-4p::hpk-1 + pCFJ90 (myo-2p::mCherry)] line 2; rmls132 [unc-54p::Q35::YFP] I* |
| AVS693 | *artEx73 [cat-2p::hpk-1 + pCFJ90 (myo-2p::mCherry)] line 1; rmls132 [unc-54p::Q35::YFP] I* |
| AVS694 | *artEx74 [cat-2p::hpk-1 + pCFJ90 (myo-2p::mCherry)] line 2; rmls132 [unc-54p::Q35::YFP] I* |
| AVS809 | *artEx62 [tph-1p::hpk-1 + pCFJ90 (myo-2p::mCherry)] line 1; sqs13 [lgg-1p::GFP::lgg-1 + odr-1p::RFP]* |
| AVS794 | *artEx65 [unc-47p::hpk-1 + pCFJ90 (myo-2p::mCherry)] line 1; sqs13 [lgg-1p::GFP::lgg-1 + odr-1p::RFP]* |
| AVS709 | *artEx62 [tph-1p::hpk-1 + pCFJ90 (myo-2p::mCherry)] line 1* |
| AVS710 | *artEx65 [unc-47p::hpk-1 + pCFJ90 (myo-2p::mCherry)] line 1* |
| AVS836 | *artEx91 [hpk-1p::GFP + tph-1p::mCherry] line 1* |
| AVS757 | *artEx87 [hpk-1p::mCherry + unc-47p::GFP] line 1* |
| AVS816 | *artEx41(rab-3p::hpk-1::CFP + pCFJ90 (myo-2p::mCherry); sqs24 [lgg-1p::GFP::lgg-1 + unc-122p::RFP]; unc-30(ok613)* |
| AVS872 | *artEx41(rab-3p::hpk-1::CFP + pCFJ90 (myo-2p::mCherry); sqs24 [lgg-1p::GFP::lgg-1 + unc-122p::RFP]; tph-1(mg280)* |

Strains with both an AVS# and an external designation were generated in another laboratory. In these cases, an AVS# is assigned for internal record keeping and increased rigor and reproducibility purposes.

**Table S2.**
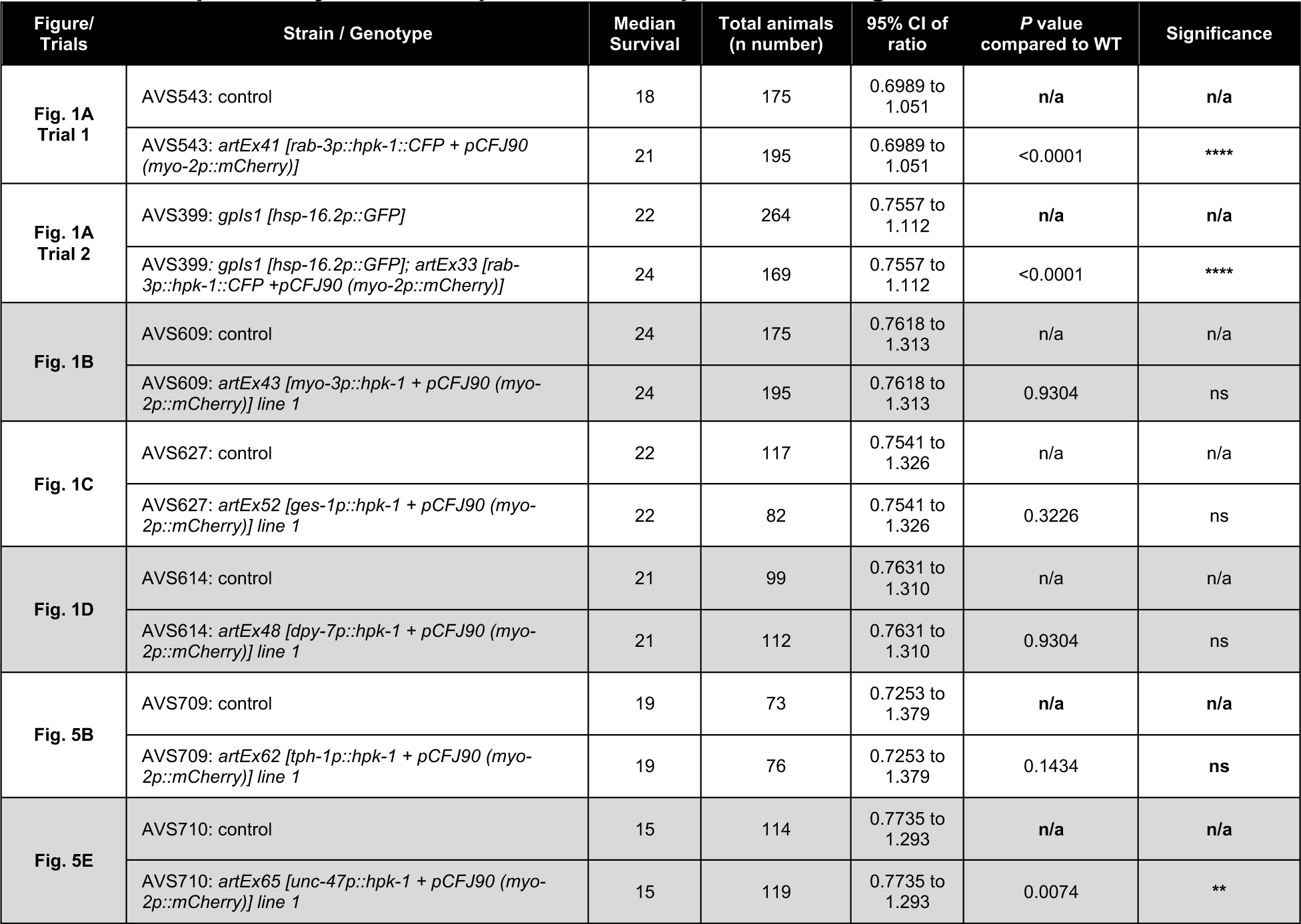
Lifespan assays data of representative experiments in figures and additional trials.

**Table S3.** Polyglutamine aggregates data of representative experiments in figures and additional trials.

| Figure/<br>Trials | Strain / Genotype | RNAi | Days of<br>adulthood | #Foci/animal<br>(mean) | Total cells<br>(n number) | SEM | P value<br>compared to<br>control | Significance |
| --- | --- | --- | --- | --- | --- | --- | --- | --- |
| **Fig. 2B**<br>**Line 1** | AVS832: *rmls132 [unc-54p::Q35::YFP] I* | n/a | 1 | 13.43 | 28 | 0.45 | n/a | n/a |
|  | AVS832: *rmls132 [unc-54p::Q35::YFP] I* | n/a | 2 | 16.62 | 21 | 0.95 | n/a | n/a |
|  | AVS832: *rmls132 [unc-54p::Q35::YFP] I* | n/a | 3 | 26 | 18 | 1.15 | n/a | n/a |
|  | AVS832: *rmls132 [unc-54p::Q35::YFP] I; artEx58 [rab-3p::hpk-1::CFP + pCFJ90 (myo-2p::mCherry)]* | n/a | 1 | 11.73 | 22 | 0.4798 | 0.013 | * |
|  | AVS832: *rmls132 [unc-54p::Q35::YFP] I; artEx58 [rab-3p::hpk-1::CFP + pCFJ90 (myo-2p::mCherry)]* | n/a | 2 | 14.11 | 19 | 0.5181 | 0.0303 | * |
|  | AVS832: *rmls132 [unc-54p::Q35::YFP] I; artEx58 [rab-3p::hpk-1::CFP + pCFJ90 (myo-2p::mCherry)]* | n/a | 3 | 21.29 | 24 | 0.9178 | 0.0024 | ** |
| **Fig. 2B**<br>**Line 2** | AVS833: *rmls132 [unc-54p::Q35::YFP] I* | n/a | 1 | 12.19 | 21 | 0.646 | n/a | n/a |
|  | AVS833: *rmls132 [unc-54p::Q35::YFP] I* | n/a | 2 | 22.18 | 11 | 1.102 | n/a | n/a |
|  | AVS833: *rmls132 [unc-54p::Q35::YFP] I* | n/a | 3 | 35 | 18 | 1.576 | n/a | n/a |
|  | AVS833: *rmls132 [unc-54p::Q35::YFP] I; artEx59 [rab-3p::hpk-1::CFP + pCFJ90 (myo-2p::mCherry)]* | n/a | 1 | 11.05 | 20 | 0.59 | 0.2017 | ns |
|  | AVS833: *rmls132 [unc-54p::Q35::YFP] I; artEx59 [rab-3p::hpk-1::CFP + pCFJ90 (myo-2p::mCherry)]* | n/a | 2 | 16.16 | 19 | 0.76 | <0.0001 | **** |
|  | AVS833: *rmls132 [unc-54p::Q35::YFP] I; artEx59 [rab-3p::hpk-1::CFP + pCFJ90 (myo-2p::mCherry)]* | n/a | 3 | 30.63 | 24 | 1.17 | 0.0282 | * |
| Fig. 2B<br>Line 3 | AVS834: *rmls132 [unc-54p::Q35::YFP] I* | n/a | 1 | 12.91 | 22 | 0.75 | n/a | n/a |
|  | AVS834: *rmls132 [unc-54p::Q35::YFP] I* | n/a | 2 | 19.83 | 23 | 0.86 | n/a | n/a |
|  | AVS834: *rmls132 [unc-54p::Q35::YFP] I* | n/a | 3 | 28 | 21 | 1.26 | n/a | n/a |
|  | AVS834: *rmls132 [unc-54p::Q35::YFP] I; artEx60 [rab-3p::hpk-1::CFP + pCFJ90 (myo-2p::mCherry)]* | n/a | 1 | 10.5 | 26 | 0.54 | 0.0105 | * |
|  | AVS834: *rmls132 [unc-54p::Q35::YFP] I; artEx60 [rab-3p::hpk-1::CFP + pCFJ90 (myo-2p::mCherry)]* | n/a | 2 | 16.12 | 26 | 0.69 | 0.0014 | ** |
|  | AVS834: *rmls132 [unc-54p::Q35::YFP] I; artEx60 [rab-3p::hpk-1::CFP + pCFJ90 (myo-2p::mCherry)]* | n/a | 3 | 20.06 | 31 | 0.81 | <0.0001 | **** |
| Fig. 2B<br>Line 4 | AVS835: *rmls132 [unc-54p::Q35::YFP] I* | n/a | 1 | 12.14 | 21 | 0.48 | n/a | n/a |
|  | AVS835: *rmls132 [unc-54p::Q35::YFP] I* | n/a | 2 | 20.78 | 23 | 0.8 | n/a | n/a |
|  | AVS835: *rmls132 [unc-54p::Q35::YFP] I* | n/a | 3 | 26.79 | 19 | 1 | n/a | n/a |
|  | AVS835: *rmls132 [unc-54p::Q35::YFP] I; artEx61 [rab-3p::hpk-1::CFP + pCFJ90 (myo-2p::mCherry)]* | n/a | 1 | 8.593 | 27 | 0.507 | <0.0001 | **** |
|  | AVS835: *rmls132 [unc-54p::Q35::YFP] I; artEx61 [rab-3p::hpk-1::CFP + pCFJ90 (myo-2p::mCherry)]* | n/a | 2 | 16.47 | 36 | 0.455 | <0.0001 | **** |
|  | AVS835: *rmls132 [unc-54p::Q35::YFP] I; artEx61 [rab-3p::hpk-1::CFP + pCFJ90 (myo-2p::mCherry)]* | n/a | 3 | 21.58 | 33 | 0.903 | 0.0005 | *** |
| Fig. 2B<br>Line 5 | AVS562: *rmls132 [unc-54p::Q35::YFP] I* | n/a | 1 | 7.185 | 27 | 0.453 | n/a | n/a |
|  | AVS562: *rmls132 [unc-54p::Q35::YFP] I* | n/a | 2 | 15.16 | 32 | 0.775 | n/a | n/a |
|  | AVS562: *rmls132 [unc-54p::Q35::YFP] I* | n/a | 3 | 28.05 | 39 | 0.99 | n/a | n/a |
|  | AVS562: *rmls132 [unc-54p::Q35::YFP] I* | n/a | 4 | 41 | 25 | 1.268 | n/a | n/a |
|  | AVS562: *rmls132 [unc-54p::Q35::YFP] I; artEx41 [rab-3p::hpk-1::CFP + pCFJ90 (myo-2p::mCherry)]* | n/a | 1 | 5.409 | 22 | 0.495 | 0.0111 | * |
|  | AVS562: *rmls132 [unc-54p::Q35::YFP] I; artEx41 [rab-3p::hpk-1::CFP + pCFJ90 (myo-2p::mCherry)]* | n/a | 2 | 11.05 | 20 | 0.956 | 0.0017 | ** |
|  | AVS562: *rmls132 [unc-54p::Q35::YFP] I; artEx41 [rab-3p::hpk-1::CFP + pCFJ90 (myo-2p::mCherry)]* | n/a | 3 | 21.14 | 21 | 0.996 | <0.0001 | **** |
|  | AVS562: *rmls132 [unc-54p::Q35::YFP] I; artEx41 [rab-3p::hpk-1::CFP + pCFJ90 (myo-2p::mCherry)]* | n/a | 4 | 32.95 | 21 | 1.702 | 0.0004 | *** |
| Fig. 2D | AVS540: *rmls132 [unc-54p::Q35::YFP] I; sid-1(pk3321)V; uIs69[unc-119p::sid-1; myo-2p::mCherry]V* | empty vector | 1 | 6.091 | 22 | 0.446 | n/a | n/a |
|  | AVS540: *rmls132 [unc-54p::Q35::YFP] I; sid-1(pk3321)V; uIs69[unc-119p::sid-1; myo-2p::mCherry]V* | empty vector | 2 | 8.125 | 24 | 0.368 | n/a | n/a |
|  | AVS540: *rmls132 [unc-54p::Q35::YFP] I; sid-1(pk3321)V; uIs69[unc-119p::sid-1; myo-2p::mCherry]V* | empty vector | 3 | 18.64 | 25 | 0.781 | n/a | n/a |
|  | AVS540: *rmls132 [unc-54p::Q35::YFP] I; sid-1(pk3321)V; uIs69[unc-119p::sid-1; myo-2p::mCherry]V* | empty vector | 4 | 32.48 | 29 | 1.105 | n/a | n/a |
|  | AVS540: *rmls132 [unc-54p::Q35::YFP] I; sid-1(pk3321)V; uIs69[unc-119p::sid-1; myo-2p::mCherry]V* | *hpk-1* | 1 | 7.609 | 23 | 0.434 | 0.0189 | * |
|  | AVS540: *rmls132 [unc-54p::Q35::YFP] I; sid-1(pk3321)V; uls69[unc-119p::sid-1; myo-2p::mCherry]V* | *hpk-1* | 2 | 12.95 | 22 | 0.64 | <0.0001 | **** |
|  | AVS540: *rmls132 [unc-54p::Q35::YFP] I; sid-1(pk3321)V; uls69[unc-119p::sid-1; myo-2p::mCherry]V* | *hpk-1* | 3 | 27.12 | 26 | 1.209 | <0.0001 | **** |
|  | AVS540: *rmls132 [unc-54p::Q35::YFP] I; sid-1(pk3321)V; uls69[unc-119p::sid-1; myo-2p::mCherry]V* | *hpk-1* | 4 | 45.32 | 28 | 1.04 | <0.0001 | **** |
| Fig. 2G,H | AVS562: *rmls132 [unc-54p::Q35::YFP] I* | n/a | 3 | 28.05 | 39 | 0.9898 | n/a | n/a |
|  | AVS562: *rmls132 [unc-54p::Q35::YFP] I; artEx41 [rab-3p::hpk-1::CFP + pCFJ90 (myo-2p::mCherry)]* | n/a | 3 | 21.14 | 21 | 0.9959 | n/a | n/a |
|  | AVS713: *rmls132 [unc-54p::Q35::YFP] I; unc-31(e169)V* | n/a | 3 | 21.52 | 58 | 0.6527 | n/a | n/a |
|  | AVS713: *rmls132 [unc-54p::Q35::YFP] I; artEx41 [rab-3p::hpk-1::CFP + pCFJ90 (myo-2p::mCherry)]; unc-31(e169)V* | n/a | 3 | 18.02 | 50 | 0.5859 | Compared to AVS713 control: 0.0001 | *** |
|  | AVS810*rmls132 [unc-54p::Q35::YFP] I; unc-13(e51)I* | n/a | 3 | 57.81 | 26 | 1.098 | n/a | n/a |
|  | AVS810*rmls132 [unc-54p::Q35::YFP] I; artEx41 [rab-3p::hpk-1::CFP + pCFJ90 (myo-2p::mCherry)]; unc-13(e51)I* | n/a | 3 | 58.68 | 25 | 1.408 | Compared to AVS810 control: 0.6258 | ns |
| Fig. 4A Line 1 | AVS682: *rmls132 [unc-54p::Q35::YFP] I* | n/a | 1 | 6.563 | 32 | 0.301 | n/a | n/a |
|  | AVS682: *rmls132 [unc-54p::Q35::YFP] I* | n/a | 2 | 17.43 | 30 | 0.7786 | n/a | n/a |
|  | AVS682: *rmls132 [unc-54p::Q35::YFP] I* | n/a | 3 | 40.23 | 26 | 1.017 | n/a | n/a |
|  | AVS682: *artEx62 [tph-1p::hpk-1 + pCFJ90 (myo-2p::mCherry)] line 1; rmls132 [unc-54p::Q35::YFP] I* | n/a | 1 | 5.333 | 21 | 0.252 | 0.0056 | ** |
|  | AVS682: *artEx62 [tph-1p::hpk-1 + pCFJ90 (myo-2p::mCherry)] line 1; rmls132 [unc-54p::Q35::YFP] I* | n/a | 2 | 15.24 | 25 | 0.609 | 0.0359 | * |
|  | AVS682: *artEx62 [tph-1p::hpk-1 + pCFJ90 (myo-2p::mCherry)] line 1; rmls132 [unc-54p::Q35::YFP] I* | n/a | 3 | 37.06 | 17 | 1.152 | 0.0494 | * |
| Fig. 4A Line 2 | AVS683: *rmls132 [unc-54p::Q35::YFP] I* | n/a | 3 | 31.43 | 21 | 1.245 | n/a | n/a |
|  | AVS683: *artEx63 [tph-1p::hpk-1 + pCFJ90 (myo-2p::mCherry)] line 1; rmls132 [unc-54p::Q35::YFP] I* | n/a | 3 | 26.29 | 24 | 1.182 | 0.0046 | ** |
| Fig. 4C Line 1 | AVS685: *rmls132 [unc-54p::Q35::YFP] I* | n/a | 1 | 6.737 | 19 | 0.3734 | n/a | n/a |
|  | AVS685: *rmls132 [unc-54p::Q35::YFP] I* | n/a | 2 | 12.26 | 19 | 0.696 | n/a | n/a |
|  | AVS685: *rmls132 [unc-54p::Q35::YFP] I* | n/a | 3 | 29.67 | 21 | 0.9394 | n/a | n/a |
|  | AVS685: *artEx65 [unc-47p::hpk-1 + pCFJ90 (myo-2p::mCherry)] line 1; rmls132 [unc-54p::Q35::YFP] I* | n/a | 1 | 6 | 25 | 0.2449 | 0.0937 | ns |
|  | AVS685: *artEx65 [unc-47p::hpk-1 + pCFJ90 (myo-2p::mCherry)] line 1; rmls132 [unc-54p::Q35::YFP] I* | n/a | 2 | 10.32 | 25 | 0.7158 | 0.064 | ns |
|  | AVS685: *artEx65 [unc-47p::hpk-1 + pCFJ90 (myo-2p::mCherry)] line 1; rmls132 [unc-54p::Q35::YFP] I* | n/a | 3 | 25.5 | 26 | 0.8931 | 0.0026 | ** |
| Fig. 4C Line 2 | AVS686: *rmls132 [unc-54p::Q35::YFP] I* | n/a | 3 | 37.32 | 22 | 1.223 | n/a | n/a |
|  | AVS686: *artEx66 [unc-47p::hpk-1 + pCFJ90 (myo-2p::mCherry)] line 1; rmls132 [unc-54p::Q35::YFP] I* | n/a | 3 | 33.32 | 22 | 1.058 | 0.0175 | * |
| Fig. S2A Line 1 | AVS691: *rmls132 [unc-54p::Q35::YFP] I* | n/a | 1 | 6.44 | 25 | 0.2242 | n/a | n/a |
|  | AVS691: *rmls132 [unc-54p::Q35::YFP] I* | n/a | 2 | 10 | 32 | 0.4283 | n/a | n/a |
|  | AVS691: *rmls132 [unc-54p::Q35::YFP] I* | n/a | 3 | 29.6 | 20 | 1.663 | n/a | n/a |
|  | AVS691: *artEx71 [eat-4p::hpk-1 + pCFJ90 (myo-2p::mCherry)] line 1; rmls132 [unc-54p::Q35::YFP] I* | n/a | 1 | 5.444 | 18 | 0.2826 | 0.0079 | ** |
|  | AVS691: *artEx71 [eat-4p::hpk-1 + pCFJ90 (myo-2p::mCherry)] line 1; rmls132 [unc-54p::Q35::YFP] I* | n/a | 2 | 10.1 | 21 | 0.5811 | 0.8935 | ns |
|  | AVS691: *artEx71 [eat-4p::hpk-1 + pCFJ90 (myo-2p::mCherry)] line 1; rmls132 [unc-54p::Q35::YFP] I* | n/a | 3 | 28.48 | 23 | 1.204 | 0.5812 | ns |
| Fig. S2A<br>Line 2 | AVS692: *rmls132 [unc-54p::Q35::YFP] I* | n/a | 3 | 23.73 | 22 | 1.273 | n/a | n/a |
|  | AVS692: *artEx72 [eat-4p::hpk-1 + pCFJ90 (myo-2p::mCherry)] line 1; rmls132 [unc-54p::Q35::YFP] I* | n/a | 3 | 22.64 | 22 | 1.118 | 0.523 | ns |
| Fig. S2C<br>Line 1 | AVS693: *rmls132 [unc-54p::Q35::YFP] I* | n/a | 1 | 6.906 | 36 | 0.1871 | n/a | n/a |
|  | AVS693: *rmls132 [unc-54p::Q35::YFP] I* | n/a | 2 | 12.77 | 26 | 0.6801 | n/a | n/a |
|  | AVS693: *rmls132 [unc-54p::Q35::YFP] I* | n/a | 3 | 32.88 | 33 | 0.7986 | n/a | n/a |
|  | AVS693: *artEx73 [cat-2p::hpk-1 + pCFJ90 (myo-2p::mCherry)] line 1; rmls132 [unc-54p::Q35::YFP] I* | n/a | 1 | 7.563 | 16 | 0.3648 | 0.0816 | ns |
|  | AVS693: *artEx73 [cat-2p::hpk-1 + pCFJ90 (myo-2p::mCherry)] line 1; rmls132 [unc-54p::Q35::YFP] I* | n/a | 2 | 12.15 | 20 | 0.8088 | 0.5587 | ns |
|  | AVS693: *artEx73 [cat-2p::hpk-1 + pCFJ90 (myo-2p::mCherry)] line 1; rmls132 [unc-54p::Q35::YFP] I* | n/a | 3 | 33.21 | 24 | 1.07 | 0.8019 | ns |
| Fig. S2C<br>Line 2 | AVS694: *rmls132 [unc-54p::Q35::YFP] I* | n/a | 3 | 22.64 | 22 | 1.118 | n/a | n/a |
|  | AVS694: *artEx74 [cat-2p::hpk-1 + pCFJ90 (myo-2p::mCherry)] line 1; rmls132 [unc-54p::Q35::YFP] I* | n/a | 3 | 23.73 | 22 | 1.273 | 0.574 | ns |

**Table S4.** Body bends assay data.

| Figure/<br>Trials | Strain / Genotype | RNAi | Body bends mean<br>(30 seconds) | Total<br>animals<br>(n number) | SEM | P value<br>compared to<br>control | Significance |
| --- | --- | --- | --- | --- | --- | --- | --- |
| Fig. 1E | N2: *wild type* | n/a | 48.72 | 40 | 0.7938 | n/a | n/a |
|  | AVS392: *hpk-1(pk1393)* | n/a | 27.35 | 41 | 1.047 | <0.0001 | **** |
|  | AVS420: *hpk-1(pk1393) X; artEx41 [rab-3p::hpk-1::CFP + pCFJ90 (myo-2p::mCherry)]* | n/a | 36.22 | 50 | 1.323 | Compared to<br>*hpk-1(pk1393)*:<br><0.0001 | **** |

**Table S5.**
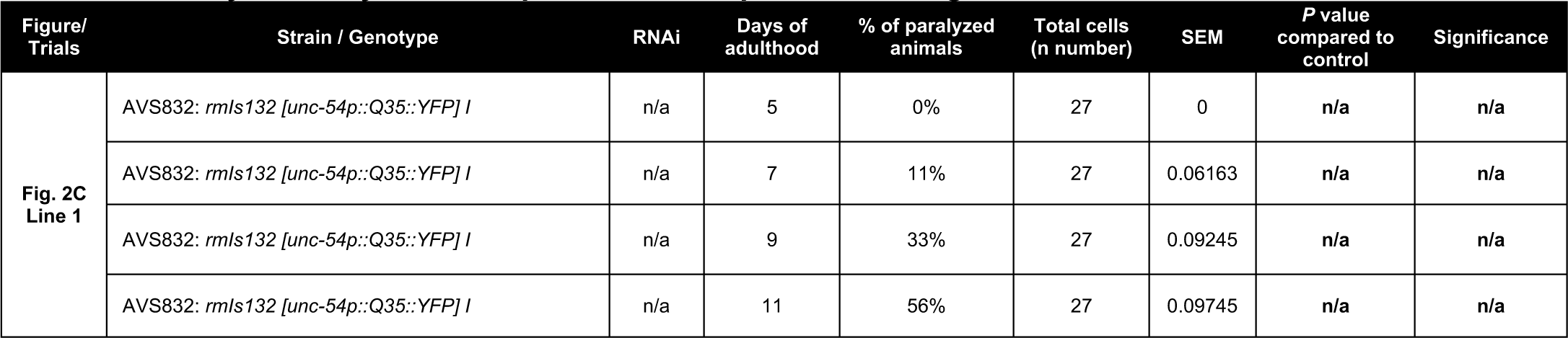
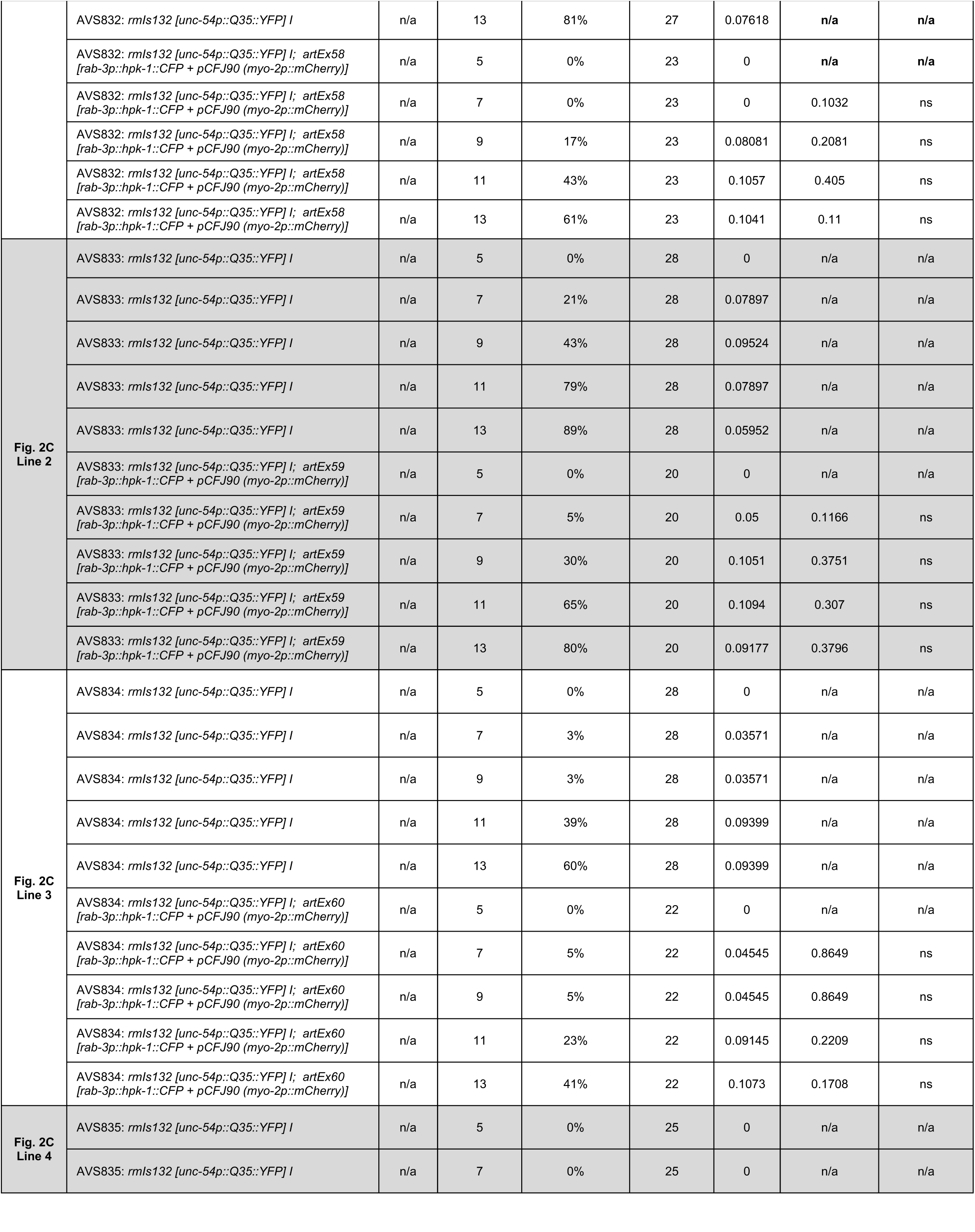
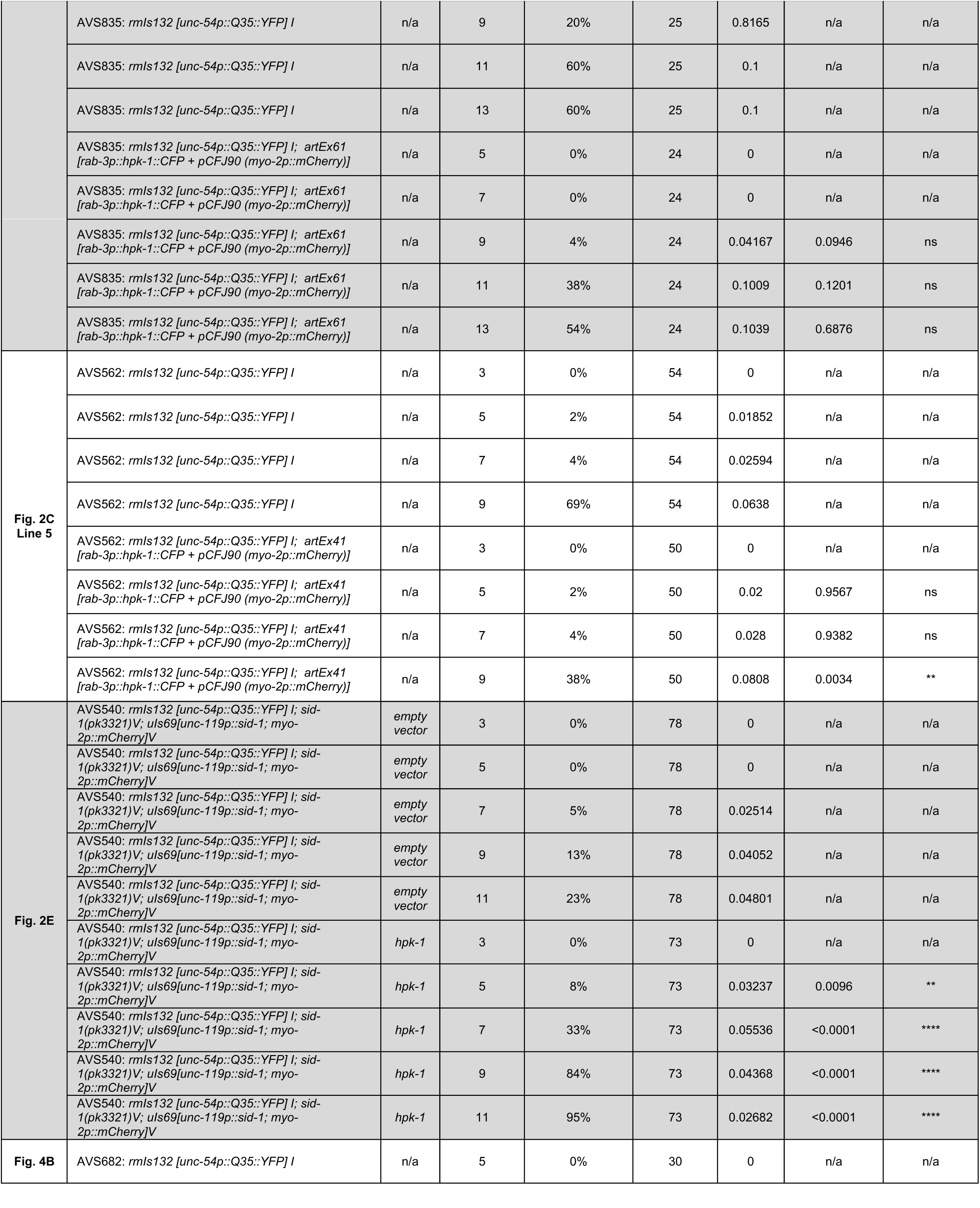
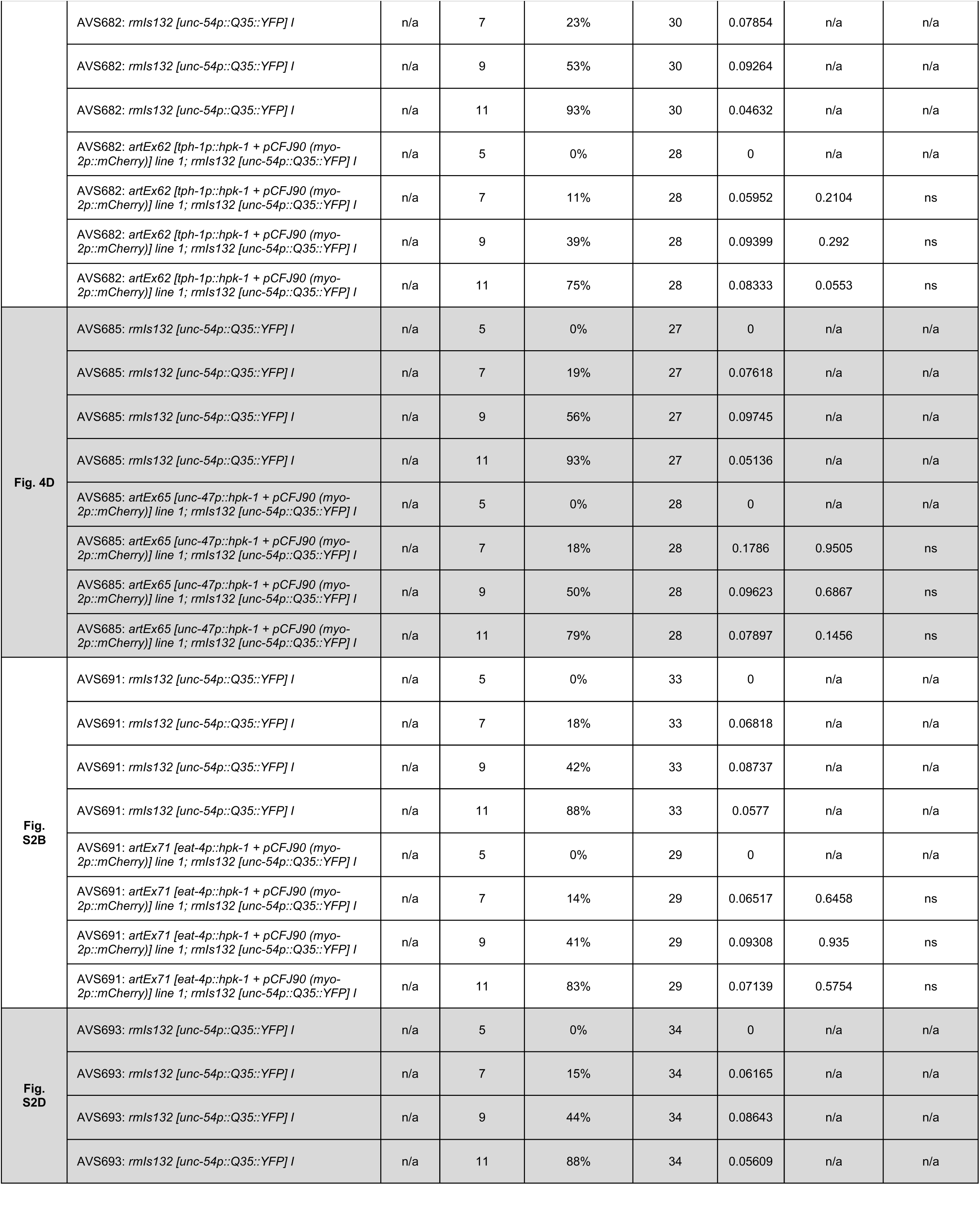
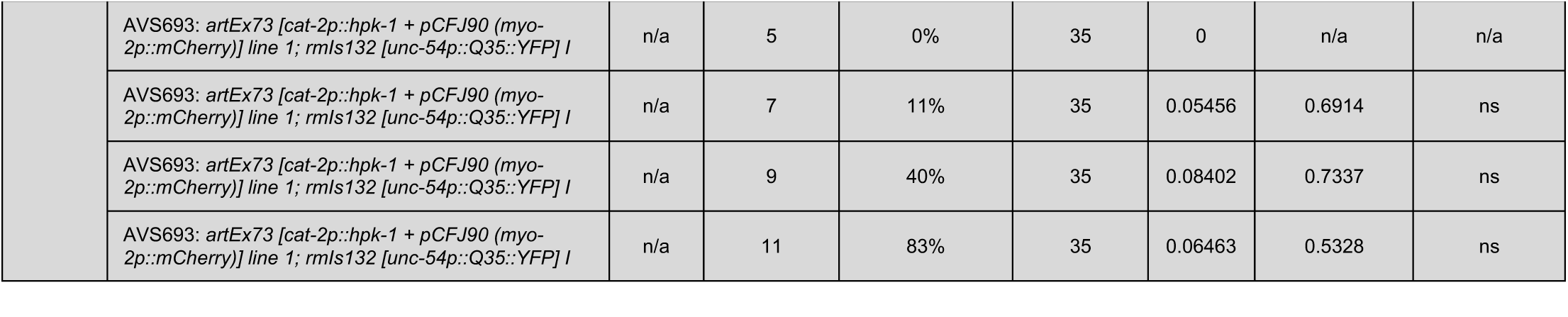
Paralysis assay data of representative experiments in figures and additional trials.

**Table S6.** Thermotolerance assay data of all representative experiments in figures and other trials.

| Figure/<br>Trials | Strain / Genotype | Exposure<br>time at<br>35C | Survival<br>% | Total animals<br>(n number) | SEM | P value<br>compared to<br>control | Significance |
| --- | --- | --- | --- | --- | --- | --- | --- |
| Fig. 1F<br>Trial 1 | N2 | 10h | 80% | 220 | 0.0268 | n/a | n/a |
|  | *hpk-1(pk1393) X* | 10h | 48% | 27 | 0.09799 | <0.0001 | **** |
|  | *hpk-1(pk1393) X; artEx41 [rab-3p::hpk-1::CFP + pCFJ90 (myo-2p::mCherry)]* | 10h | 82% | 50 | 0.05488 | Compared to<br>*hpk-1(pk1393)*:<br>0.0016 | ** |
| Fig. 1F<br>Trial 2 | N2 | 10h | 64% | 130 | 0.00251 | n/a | n/a |
|  | *hpk-1(pk1393) X* | 10h | 27% | 66 | 0.01088 | <0.0001 | **** |
|  | *hpk-1(pk1393) X; artEx41 [rab-3p::hpk-1::CFP + pCFJ90 (myo-2p::mCherry)]* | 10h | 76% | 75 | 0.00965 | Compared to<br>*hpk-1(pk1393)*:<br><0.0001 | **** |
| Fig. 5A | AVS709: control | 6h | 84% | 32 | 0.06521 | n/a | n/a |
|  | AVS709: control | 8h | 67% | 39 | 0.07647 | n/a | n/a |
|  | AVS709: *artEx62 [tph-1p::hpk-1 + pCFJ90 (myo-2p::mCherry)] line 1* | 6h | 100% | 33 | 0 | 0.0178 | * |
|  | AVS709: *artEx62 [tph-1p::hpk-1 + pCFJ90 (myo-2p::mCherry)] line 1* | 8h | 83% | 41 | 0.05949 | 0.0955 | ns |
| Fig. 5D | AVS710: control | 6h | 85% | 54 | 0.0488 | n/a | n/a |
|  | AVS710: control | 8h | 69% | 49 | 0.6652 | n/a | n/a |
|  | AVS710: *artEx65 [unc-47p::hpk-1 + pCFJ90 (myo-2p::mCherry)] line 1* | 6h | 89% | 47 | 0.04546 | 0.5366 | ns |
|  | AVS710: *artEx65 [unc-47p::hpk-1 + pCFJ90 (myo-2p::mCherry)] line 1* | 8h | 69% | 51 | 0.06562 | 0.9353 | ns |

**Table S7.**
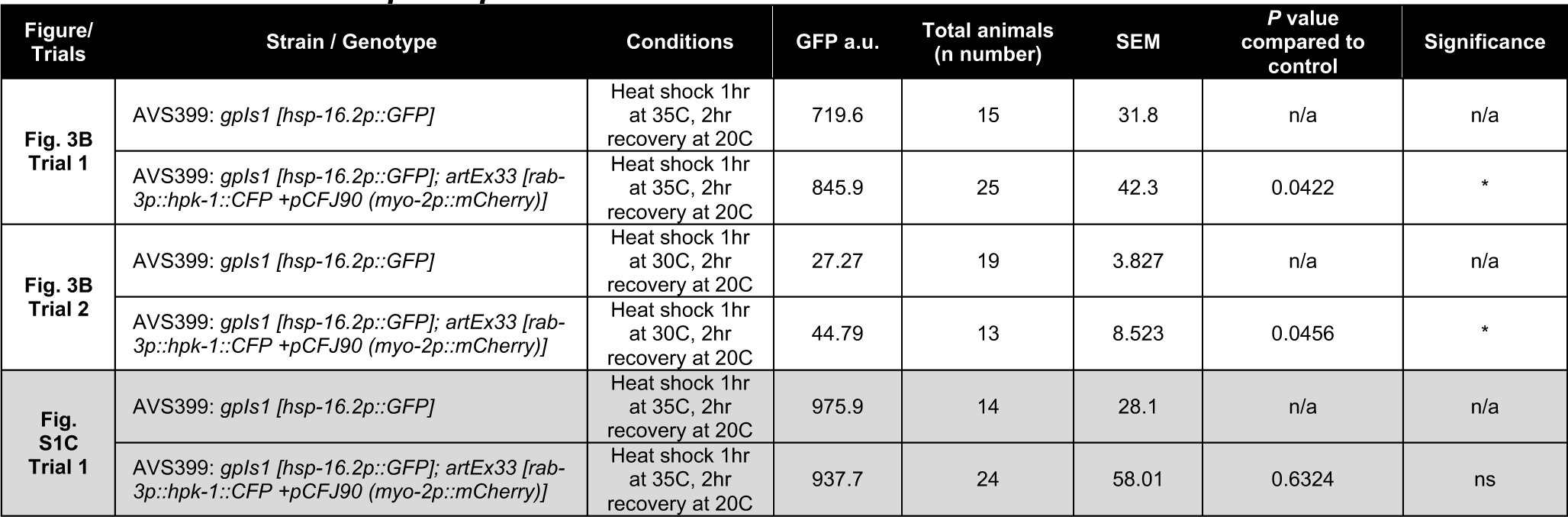
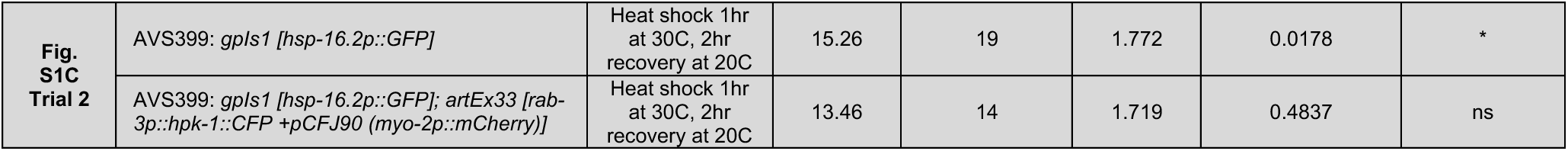
Induction of *hsp-16.2p::GFP* data after heat stress.

**Table S8.** Induction of autophagosomes at basal conditions data.

| Figure/<br>Trials | Strain / Genotype | Age | # of<br>autophagosomes | Total cells<br>(n number) | SEM | P value<br>compared to<br>control | Significance |
| --- | --- | --- | --- | --- | --- | --- | --- |
| **Fig. 3D**<br>**Trial 1** | AVS716: *sqli13 [lgg-1p::GFP::lgg-1 + odr-1p::RFP]* | L4 | 1.576 | 59 | 0.1214 | n/a | n/a |
|  | AVS716: *artEx41 [rab-3p::hpk-1::CFP + pCFJ90 (myo-2p::mCherry)]; sqli13 [lgg-1p::GFP::lgg-1 + odr-1p::RFP]* | L4 | 2.651 | 43 | 0.166 | <0.0001 | **** |
| **Fig. 3D**<br>**Trial 2** | AVS716: *sqli13 [lgg-1p::GFP::lgg-1 + odr-1p::RFP]* | L4 | 1.863 | 80 | 0.1107 | n/a | n/a |
|  | AVS716: *sqli13 [lgg-1p::GFP::lgg-1 + odr-1p::RFP]* | D1 | 2.597 | 67 | 0.1295 | n/a | n/a |
|  | AVS716: *artEx41 [rab-3p::hpk-1::CFP + pCFJ90 (myo-2p::mCherry)]; sqli13 [lgg-1p::GFP::lgg-1 + odr-1p::RFP]* | L4 | 2.418 | 79 | 0.1355 | 0.0018 | ** |
|  | AVS716: *artEx41 [rab-3p::hpk-1::CFP + pCFJ90 (myo-2p::mCherry)]; sqli13 [lgg-1p::GFP::lgg-1 + odr-1p::RFP]* | D1 | 3.574 | 61 | 0.1661 | <0.0001 | **** |
| **Fig. 5C** | AVS809: *sqli13 [lgg-1p::GFP::lgg-1 + odr-1p::RFP]* | D1 | 1.238 | 80 | 0.09285 | n/a | n/a |
|  | AVS809: *artEx62 [tph-1p::hpk-1 + pCFJ90 (myo-2p::mCherry)] line 1; sqli13 [lgg-1p::GFP::lgg-1 + odr-1p::RFP]* | D1 | 1.356 | 104 | 0.08074 | 0.3372 | ns |
| **Fig. 5F** | AVS794: *sqli13 [lgg-1p::GFP::lgg-1 + odr-1p::RFP]* | D1 | 1.672 | 64 | 0.1221 | n/a | n/a |
|  | AVS794: *artEx65 [unc-47p::hpk-1 + pCFJ90 (myo-2p::mCherry)] line 1; sqli13 [lgg-1p::GFP::lgg-1 + odr-1p::RFP]* | D1 | 2.329 | 73 | 0.1051 | <0.0001 | **** |
| **Fig. S1E**<br>**Trial 1** | AVS716: *sqli13 [lgg-1p::GFP::lgg-1 + odr-1p::RFP]* | L4 | 21 | 11 | 2.347 | n/a | n/a |
|  | AVS716: *artEx41 [rab-3p::hpk-1::CFP + pCFJ90 (myo-2p::mCherry)]; sqli13 [lgg-1p::GFP::lgg-1 + odr-1p::RFP]* | L4 | 19 | 11 | 2.106 | 0.5331 | ns |
| **Fig. S1E**<br>**Trial 2** | AVS716: *sqli13 [lgg-1p::GFP::lgg-1 + odr-1p::RFP]* | L4 | 23.57 | 23 | 2.51 | n/a | n/a |
|  | AVS716: *artEx41 [rab-3p::hpk-1::CFP + pCFJ90 (myo-2p::mCherry)]; sqli13 [lgg-1p::GFP::lgg-1 + odr-1p::RFP]* | L4 | 21.35 | 17 | 3.313 | 0.5905 | ns |
| **Fig. 5G** | AVS872: *sqli24 [lgg-1p::GFP::lgg-1 + unc-122p::RFP]; tph-1(mg280)* | D1 | 1 | 73 | 0.1068 | n/a | n/a |
|  | AVS872: *artEx41(rab-3p::hpk-1::CFP + pCFJ90 (myo-2p::mCherry)); sqli24 [lgg-1p::GFP::lgg-1 + unc-122p::RFP]; tph-1(mg280)* | D1 | 2.04 | 67 | 0.1465 | <0.0001 | **** |
|  | AVS816: *sqli24 [lgg-1p::GFP::lgg-1 + unc-122p::RFP]; unc-30(ok613)* | D1 | 1.36 | 75 | 0.1033 | n/a | n/a |
|  | AVS816: *artEx41(rab-3p::hpk-1::CFP + pCFJ90 (myo-2p::mCherry)); sqli24 [lgg-1p::GFP::lgg-1 + unc-122p::RFP]; unc-30(ok613)* | D1 | 1.35 | 83 | 0.1229 | 0.9480 | ns |

